# The unseen invaders: tracking phylogeographic dynamics and genetic diversity of cryptic *Pomacea canaliculata and P. maculata* (Golden Apple Snails) across Taiwan

**DOI:** 10.1101/2024.01.20.576472

**Authors:** Pritam Banerjee, Gobinda Dey, Jyoti Prakash Maity, Kathryn A. Stewart, Raju Kumar Sharma, Michael W.Y. Chan, Kuan Hsien Lee, Chien-Yen Chen

## Abstract

The cryptic invasion of golden apple snails (*Pomacea canaliculata and P. maculata*) in Taiwan has caused significant ecological and economical damage over last few decades, however, their management remains difficult due to inadequate taxonomic identification, complex phylogeny and limited population genetic information. We aim to understand the current distribution, putative population of origin, genetic diversity and potential path of cryptic invasion of *Pomacea canaliculata and P. maculata* across Taiwan to aid in improved mitigation approaches. The present investigation conducted a nationwide survey with 254 samples collected from 41 locations from 14 counties or cities across Taiwan. We identified *P. canaliculata* and *P. maculata* based on mitochondrial COI and compared their genetic diversity across Taiwan, as well as other introduced and native countries (based on publicly available COI data) to understand the possible paths of invasion in Taiwan. Based on mitochondrial COI barcoding, sympatric and heterogeneous distributions of invasive *P. canaliculata* and *P. maculata* were noted. Our haplotype analysis and mismatch distribution suggested multiple introductions of *P. canaliculata* in Taiwan was likely originated directly from Argentina, whereas *P. maculata* was probably introduced from a single, or a few, introduction event(s) from Argentina and Brazil. Our population genetic data further demonstrated a higher haplotype and genetic diversity for *P. canaliculata* and *P. maculata* in Taiwan compared to other introduced regions. Based on our current understanding, the establishment of *P. canaliculata* and *P. maculata* is alarming and widespread beyond geopolitical borders, requiring a concerted and expedited national and international invasive species mitigation program.

## INTRODUCTION

Invasive alien species are major drivers of biodiversity loss and have massive impact on the economic status of invaded countries (Linders et al., 2019; McNeely, 2001), even considered the second most economically destructive event after natural hazards such as earthquakes or floods (Turbelin et al., 2023). Making matters worse, morphologically similar species may invade together, causing a masking effect to species distribution and establishment, further hindering effective management practices (Saltonstall et al., 2002). Without in-depth knowledge about the origins and population genetics of invasive species, management of natural landscapes and aquascapes is rendered speculative (Sakai et al., 2001).

*Pomacea canaliculata* native to South America, have been both intentionally and unintentionally introduced in several countries (Lowe et al., 2000; Zhao et al., 2022), recently becoming a serious threat to agricultural and economic development worldwide (e.g., Asia, Europe, Africa, North America, and the Pacific Islands) (Ranamukhaarachchi & Wickramasinghe, 2006; Hayes et al., 2008; Buddie et al., 2021). Several other species of *Pomacea* (e.g., *P. maculata*, *P. diffusa*, *P. scalaris*) have also been introduced across numerous countries, and among the introduced species of *Pomacea*, *P. canaliculata* and *P. maculata* are the most destructive and morphologically cryptic (Rama Rao et al., 2018) causing frequent misidentifications. This can then lead to false biodiversity information, potentially obstructing effective invasion management (Cowie et al., 2006). Molecular method implementation has recently been found to be the best option for distinguishing *P. canaliculata* and *P. maculata* (Matsukura & Wada, 2017; Banerjee et al., 2022), with population genetics analysis recently conducted to comprehend the invasive origin and distribution of *P. canaliculata* and *P. maculata* across Asia (Hayes et al., 2008; Yang et al., 2018; Dumidae et al., 2021; Yang et al., 2022; Zhao et al., 2022; Liu et al., 2023)

Taiwan encompasses a high percentage of the world’s biodiversity compared to its landmass and houses an extraordinary array of fauna that comprises evolutionary lineages from both Palearctic and Indomalaya biogeographic realms (He et al., 2018; Päckert et al, 2009). However, as an island, Taiwan is highly sensitive to invasive species (Lee et al., 2019). Indeed, several invasive alien species have been introduced to Taiwan which has negatively affected its local biodiversity. Specifically, the introduction and establishment of *P. canaliculata* in Taiwan have resulted in serious ecological and economical destruction.

Historically, Taiwan was noted as the first Asian region to import *Pomacea* spp. for commercial utilization (dietary protein supplementation and aquarium trades) during 1979-1982 (Joshi & Sebastian, 2003; Yang et al, 2006; Cheng and Kao, 2006; Wu et al, 2010). Subsequently, *Pomacea* spp. were introduced into other Asian countries for economic trade (food and aquarium trades). Initially in Taiwan, *Pomacea* spp. were considered a healthy diet alternative that contained high protein, minerals, and vitamins, thus motivating farmers in its cultivation and trade (Cheng and Kao, 2006; Hayes et al., 2008). However, the trade of *Pomacea* spp. was soon terminated because of the snail’s poor texture and high production cost. Exacerbating *Pomacea* spp. invasion, farmers were not content with the low market value and thereafter released many *Pomacea* spp. into the neighboring countryside (Cheng and Kao, 2006; Chiu et al., 2014). Almost immediately after release, *Pomacea* spp. became a serious threat to the local agricultural system possibly as a result of its high reproduction rate, tolerance to various environmental stressors, and voracious appetite (Joshi & Sebastian, 2003). The problem with *Pomacea* was first recognized in 1982, raising national-level attention due to numerous damages to local agronomy; *Pomacea* spp. spread into various agricultural lands, as well as natural wetlands impacting cultivated plants such as rice seedlings, taro, and lotus, even destroying several cultivation fields (Cheng & Kao, 2006). Making matters worse, *Pomacea* spp. acts as a vector for various zoonotic diseases (e.g., *Angiostrongylus cantonensis*). Several cases of human eosinophilic meningitis have been reported in Taiwan through either direct (consumption of raw snail meat) or indirect (contaminated food) involvement of *Pomacea* spp. (Tsai et al., 2001). More specifically, enormous annual economic losses were reported due to these invasive gastropods (Globally: 3.94 billion USD; in Asia: 3.71 billion since 1961; (Jiang et al., 2022)), with Asian countries as the predominantly affected region (Jiang et al., 2022). Across all reported gastropods, *Pomacea* spp. (particularly *P. canaliculata*) were noted to be the major contributor. Despite this, little attention has thus far been paid to understanding the invasive biology and population genetics of *P. canaliculata* in Taiwan (Cheng & Kao, 2006; Banerjee et al., 2022), which restricted the understanding of their regional (Taiwan) and potentially global spread (Sakai et al., 2001).

Originally, two species of *Pomacea, P. canaliculata* and *P. scalaris,* were reported from Taiwan, where *P. canaliculata* was noted as a destructive, dominant species and distributed nationwide. Conversely, *P. scalaris* was reported to be less invasive and limited in distribution to few sampling locations in Southern Taiwan (Cheng et al., 2006; Yang et al., 2008; Wu et al., 2010). However, a more recent molecular study revealed the presence of another species, *P. macualta*, hiding with its cryptic relative *P. canaliculata* in Southern Taiwan (Banerjee et al., 2022). They reported that misidentification is very common between *P. canaliculata* and *P. macualta* based on morphological observation alone (Banerjee et al., 2022), suggesting limited knowledge on the true distribution and invasion history of *P. maculata.* In fact, out of the three *Pomacea* species currently in Taiwan, *P. scalaris* is considered less destructive due to its smaller hatching size and inferior growth performance (Wu et al., 2010), and thus assumed geographically restricted mostly in the South of the country (Wu et al., 2011). On the other hand, *P. canaliculata* and *P. maculata* are highly invasive and eco-economically destructive in nature, yet their actual distribution remains unknown across Taiwan (Banerjee et al., 2022).

Here, our study aims to generate accurate identification of cryptic invasive *P. canaliculata* and *P. macualta*, including their nation-wide distribution and genetic diversity to better aid in adaptive management strategies. By doing so, we aspire to elucidate their (i) possible origin, thereby minimizing ongoing propagation events, and (ii) proper distribution and putative invasion path within Taiwan to maximize future eradication measures. As subsequent introductions of *P. canaliculata* and *P. maculata* may occur from trades between Taiwan into other Asian countries (e.g., China, Japan, Thailand, Malaysia) (Joshi & Sebastian, 2003), the current lack of molecular data makes global invasive management efforts especially difficult. Here too our study may expand knowledge of *Pomacea* distributions in Taiwan. To accomplish these aims, our survey conducted collections across 41 different sampling locations within 14 counties or cities across Taiwan to reveal the origin, distribution, and population genetic structure of *P. canaliculata* and *P. maculata*. We then compared the genetic diversity of *P. canaliculata* and *P. maculata* with other countries (invaded or native) to understand their possible paths of invasion.

## 2. METHODS

### 2.1 Snail sample collection and preservation

In Taiwan, *Pomacea* spp. were presumably introduced through human activities and later distributed nationwide. Since this distribution may be affected by agricultural land or other (human-induced) distribution barriers (e.g., roads, mountains), sampling locations (populations) were centered around or near cities to understand the current nationwide distribution. We then further subdivided Taiwan into different regional parts coordinates (e.g., Southern, Northern, Eastern, Western and Middle) to more generally understand the current distribution of *Pomacea* for downstream management decisions. To accomplish this, distribution of *Pomacea* spp. were surveyed across 41 locations across 14 counties or cities in Taiwan, including Hualien (HU) and Taitung (TT) from the East, Yunlin (YL), Changhua (CH), Taichung (TC), and Miaoli (MIO) from the Middle of island, Hsinchu (HS), Taoyuan (TY), Taipei (TP), and Yilan (YI) from the North, and Pingtung (PT), Kaohsiung (KSH), Tainan (TN), and Chiayi (CY) from the South. This was achieved via manual hand-collection from crop fields and drainage ditches from April 2020 to January 2023. Sampling locations and the number of samples are shown in Figure 1 and APPENDIX 1. Collected samples were transported directly to the laboratory as soon as possible (as per sampling protocols). In cases of remote sampling or locations away from the laboratory, specimens were heat-shocked using hot water (100° C) for 10-15 minutes and subsequently kept in an ice box in separate sampling bags to avoid contamination. Once in the laboratory, samples were washed thoroughly to remove soil and debris, and then if not previously applied, were heat-shocked using a microwave to remove inner tissue from shells. After removing the inner tissue from the shell, 20-50 mg of foot tissue was used to extract DNA to avoid parasite contamination, and the rest of the tissue was stored in 95% ethanol.

**Figure 1.**
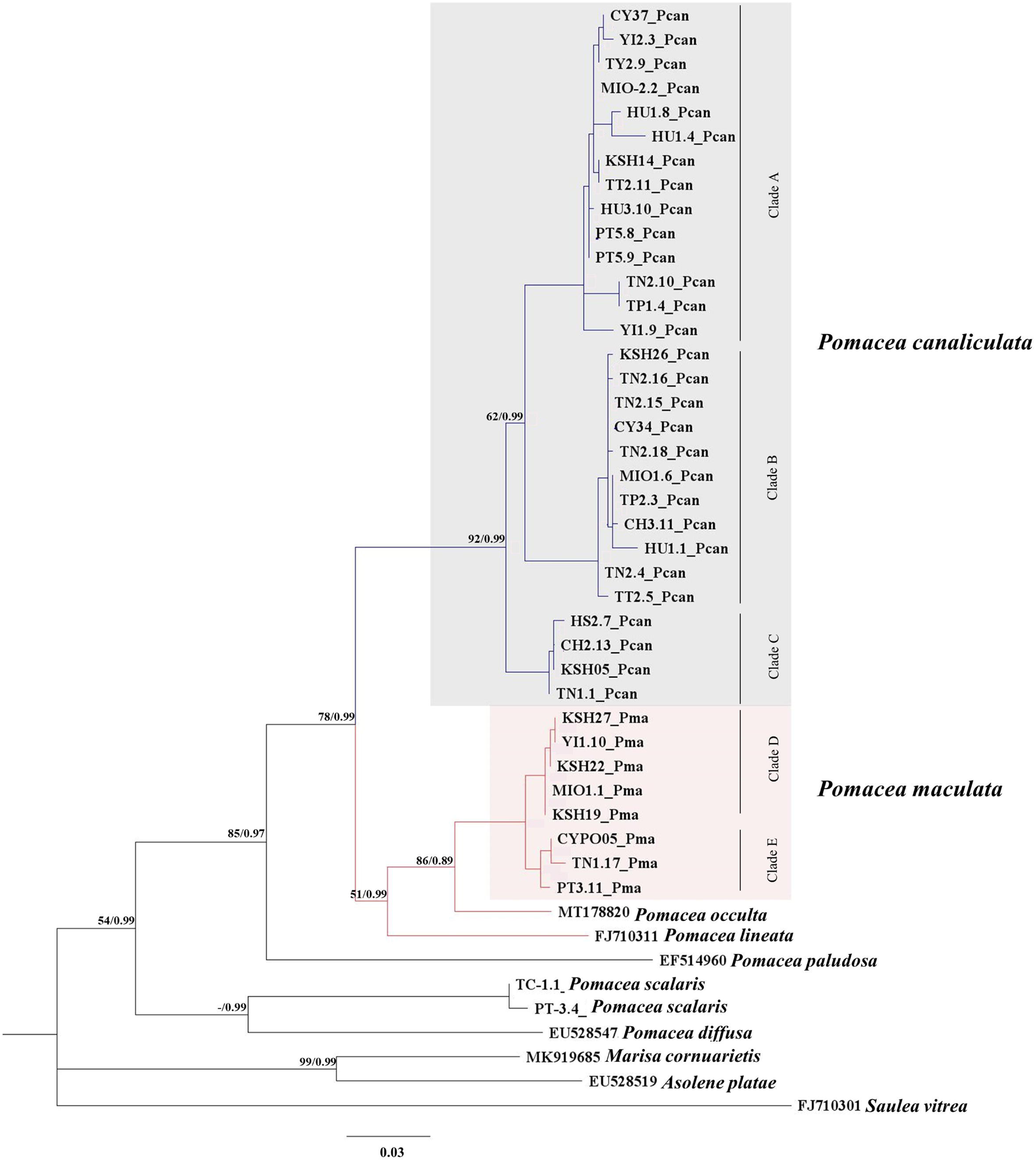
Geographical distribution and frequency haplotypes of *P. canaliculata* (Pc_Hap 1-18) and *P. maculata* (Pm_Hap 19-22) in Taiwan (Map created using ArcGIS v10.7 software)

### 2.2 Genomic DNA extraction and amplification

Genomic DNA was extracted from the 20-50 mg foot tissues of *Pomacea* spp. with the help of a DNeasy Blood and Tissue Kits (Qiagen, Germany), similar to Banerjee et al. (2022). The mitochondrial DNA barcode of *Pomacea* spp. was amplified using a conventional PCR (CLUBIO PCR system, 96-well, CB series; Scientific Biotech Corp., Taiwan) with species-specific primers: PcanCOI (F) (5’-TGG GGT ATG ATC AGG CC-3’) and PinsCOI (F) (5’-ATC TGC TGC TGT TGA AAG-3’) and HCO2198 (R) (5’-TAA ACT TCA GGG TGA CCA AAA AAT CA-3’) for initial identification of *P. canaliculata* and *P. maculata* (Matsukura et al, 2008). The double distilled water and DNA from Giant African land snail (*Lissachatina fulica*) were used as the negative control followed by the previous study (Banerjee et al., 2022). Furthermore, successfully identified specimens from initial screening were amplified using universal mitochondrial COI primers: LCO1490 (5’ GGT CAA CAA ATC ATA AAG ATA TTG G-3’) and HCO2198 (5’ TAA ACT TCA GGG TGA CAA AAA AAT CA-3’) for a 658bp fragment. The final PCR reaction contained 5 µL 5x Fast-RunTM Taq Master Mix with Dye (Protech Technology Enterprise, Taiwan), 0.5 µL of 10 pmol forward and reverse primers (Genomics, Taiwan), 1 μL DNA template from the tissue sample, for a final volume of 25 µL using ultrapure water (Thermo Fisher, USA). The PCR conditions were 5 min at 94°C, 35 cycles of 30s at 94°C (denaturation), 30s at 55°C (annealing), and 1 min at 72°C (extension), then the final extension was at 72°C for 5 min. All amplified products were sent to Genomics Co., Ltd, Taiwan for sequencing in both directions. All sequences derived from the present study were aligned using the MUSCLE algorithm (Edgar, 2004), and edited in Mega v11 (Tamura et al, 2021), and Geneious v11.1.5 (Kearse et al., 2012). A total of 254 of *Pomacea* spp. were amplified from 41 sampling spots in Taiwan.

### 2.3 Mitochondrial COI

The identification of 254 *Pomacea* spp. were accomplished via sequence similarity based on the NCBI database (https://blast.ncbi.nlm.nih.gov/Blast.cgi) with the highest level of query cover and percentage identity. To clarify the origins of our *Pomacea* spp. samples across Taiwan, previously identified and uploaded sequences of *P. canaliculata* and *P. maculata* were downloaded from NCBI which represent different geographic locations (Hayes et al., 2008; Kannan et al., 2021; Liu et al, 2019; Lv et al., 2013; Yang et al., 2018). Sequence quality was examined in Geneious v11.1.5 (Kearse et al., 2012), BioEdit v7.2 (Hall, 1999), and DnaSP v6.12.03 (Rozas et al., 2017) and filtering sequences published after 2007 to clearly distinguish *P. maculata* from *P. canaliculata* (Yang et al., 2018). A total of 1173 (948 *P. canaliculata* and 225 *P. maculata*) sequences were selected for final evaluation (APPENDIX 2, 3), and further screening was performed after a test run of phylogenetic and haplotyping network analysis (details in sections 2.5). Sequence length variation was adjusted and trimmed manually in Mega v11 (Tamura et al., 2021), allowing us to prepare three datasets, (i) the sequences generated from the present study (samples only from Taiwan) represented 611bp for *P. maculata* and *P. canaliculata* as Dataset 1, and global sequences (Taiwan and other previously published sequences collected from NCBI and/or Hayes et al., 2008; Kannan et al., 2021; Liu et al, 2019; Lv et al., 2013; Yang et al., 2018) datasets represent (ii) 579bp for *P. maculata* as Dataset 2, and (iii) 569bp for *P. canaliculata* as Dataset 3.

### 2.4 DNA polymorphism and phylogeographic distribution

All sequences were aligned in Mega v11 (Tamura et al., 2021) using the MUSCLE algorithm (Edgar, 2004), and DnaSP v6.12.03 (Rozas et al., 2017) was used to calculate the number of haplotypes (*H*) and polymorphic sites (*S*) as well as haplotype diversity (*Hd*), nucleotide diversities (π) and average number of difference (*K*). Neutrality tests such as Tajima’s *D* and Fu’s *Fs* (Fu, 1997) were performed to understand signatures of population expansion using Arlequin v3.5.2.2 (Excoffier & Lischer, 2010). Furthermore, to understand the level of genetic difference between the populations, pairwise *F_st_* values were calculated using Arlequin v3.5.2.2 (Excoffier & Lischer, 2010), and the level of gene flow/effective migration rate was calculated using the following formula: *Nm*=((1/*F_ST_*) -1)/2) (Hudson et al., 1992). The connections between haplotypes and their phylogeographic distribution were analyzed using the median-joining network implemented in PopArt v1.7 (Leigh & Bryant, 2015). Furthermore, the mismatch distribution analyses were performed in DnaSP v6.12.03 (Rozas et al., 2017) to understand the genetic variation for the population of *P. canaliculata* and *P. maculata*. The inter- and intra-specific genetic distance (*p*-distance) of *P. canaliculata* and *P. maculata* were calculated using Mega v11 (Tamura et al., 2021).

### 2.5 Phylogenetic analysis

The evolutionary model was first tested in jModelTest v2.1.10 for each population (Darriba et al., 2012). The maximum-likelihood (ML) method with the GTR+G model was used based on 1000 bootstrap replicates (Felsenstein, 1985) to build phylogenetic relations in Mega v11 (Tamura et al., 2021). A phylogenetic tree using Bayesian inference (BI) was produced using BEAUti v1.10.4 and BEAST v1.10.4 (Bouckaert et al., 2014) under GTR+G substitution model, strict clock, constant tree prior with random starting tree model. A Markov Chain Monte Carlo Analysis (MCMC) was run for 10 million generations and sampled in every 1000 generation as well as the final consensus tree was generated in TreeAnnotator v1.10.4 with 10% burn-in. The convergence states were visualized using Tracer v1.7.2 (Rambaut et al., 2018) with all the ESS values above 220. Both the maximum-likelihood (ML) and Bayesian inference (BI) analysis were run twice independently. The resulting consensus trees were visualized in FigTree v1.4.4 (Rambaut & Drummond, 2010). In all phylogenetic trees (ML and BI), *P. occulta* (FJ710311), *P. lineata* (FJ710311), *P. paludosa* (EF514960), *P. diffusa* (EU528547), *P. scalaris* (collected from TC by our team), *Marisa cornuarietis* (MK919685), *Asolene platae* (EU528547), and *Saulea vitrea* (FJ710301) were added as outgroups.

## 3. RESULTS

### 3.1 Species distribution

We used a quick identification primer (PanCOI, PinsCOI and HCO2198) and full former region (LCO1490 and HCO2198), and both identified *Pomacea* spp up to species level. Based on our collections across Taiwan, the distribution of *P. canaliculata* and *P. maculata* was observed to be sympatric, and heterogeneous. *P. canaliculata* was distributed throughout Taiwan and represented the dominant species (≈ 97%), whereas *P. maculata* represented ≈ 3% of the total number of collected individuals (Fig. 1; Map created using “ArcGIS v10.7 software; ESRI Inc., Redlands, CA, USA”). The genetic distance (p-distance) within *P. canaliculata* and *P. maculata* were 0.02 and 0.01 respectively, and genetic distance (p-distance) in between them were 0.0928 (APPENDIX 4)

### 3.2 Phylogenic analysis

Phylogenetic relationships were congruent among four independent analysis (both ML and BI) with *P. canaliculata* and *P. maculata* clustered into two clades separately (Fig. 2). *P. maculata* grouped with *Pomacea occulta* and *Pomacea lineata* together and formed a sister group of *P. canaliculata.* although this separation was not strongly supported with Maximum likelihood analysis (<80, Fig. 2). Furthermore, there were three clades noted for *P. canaliculata* (Clade A, B, and C) and two clades for *P. maculata* (Clade D and E).

**Figure 2.**
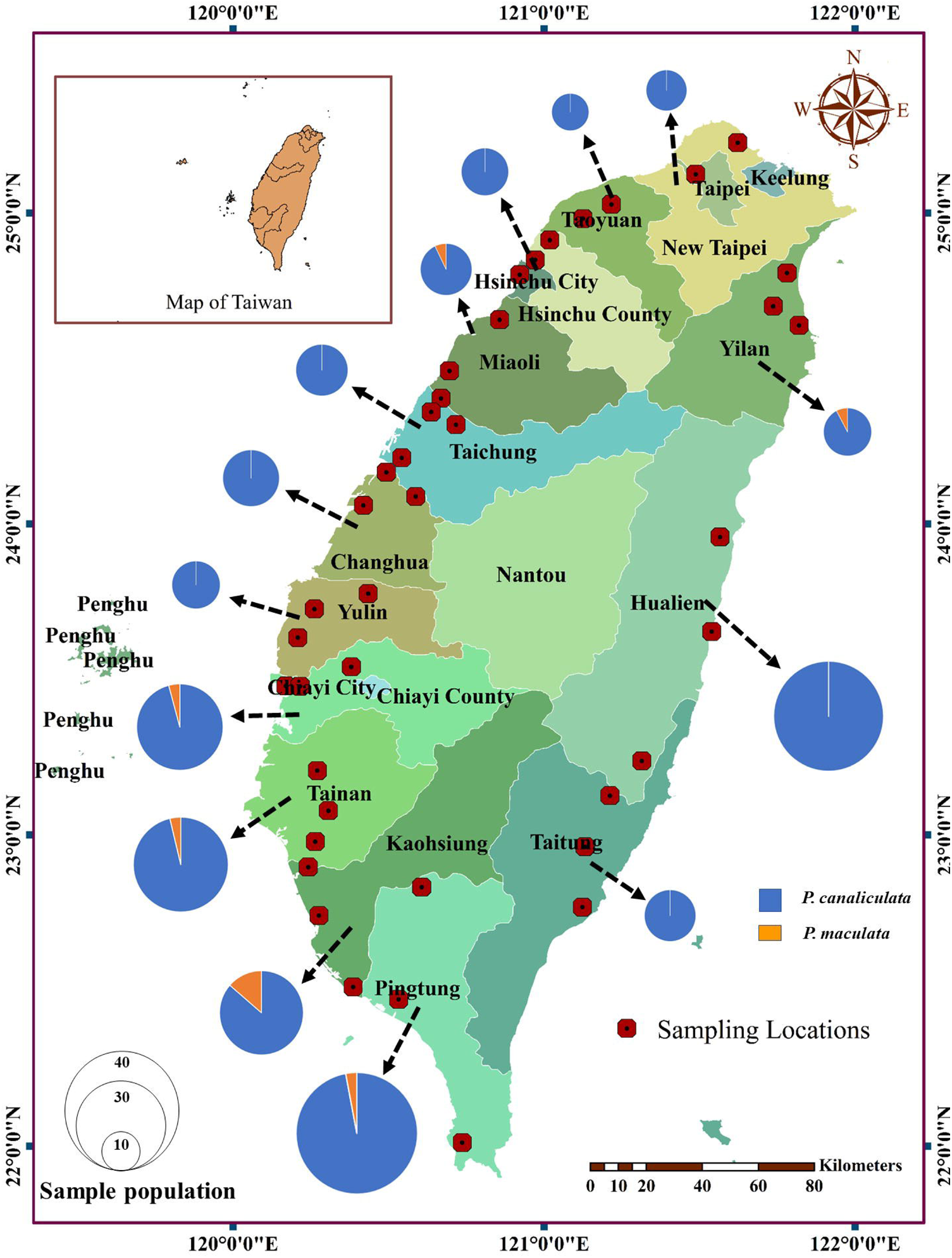
Phylogenetic tree generated from 254 sequences mitochondrial COI sequence of *P. canaliculata* and *P. maculata* collected from 41 different location (14 counties/cities) in Taiwan. Node labels are maximum likelihood/Bayesian posterior probabilities. The sequences of *P. lineata*, *P. paludosa, P. diffusa*, *P. scalaris*, *Marisa cornuarietis*, *Asolene platae*, *Saulea vitrea* were used as outgroups.

### 3.3 Haplotype distribution and network analysis

#### 3.3.1 Phylogeography of *P. canaliculata* in Taiwan

A total number of 246 sequences of *P. canaliculata* from 14 different cities/counties revealed the presence of 18 haplotypes (Pc_Hap1-18) and three different networks (Network Tw-A-C) across Taiwan (Fig. 3, Table 1, 2, APPENDIX 5). The first network (Network Tw-A) was most dominant, representing 69.90% of all sequences with seven haplotypes. Among the seven haplotypes, Pc_Hap 3 was widespread and dominant; the other six haplotypes included five unique haplotypes from Tainan (Pc_Hap 1, 6), Taitung (Pc_Hap 16), Chiayi (Pc_Hap 12), Changhua (Pc_Hap 15), and one shared by Chiayi, Yilan, Changhua, and Taichung (Pc_Hap 11). The Pc_Hap 3 demonstrated the highest number of connections which created a star-like appearance, could be a potential founder haplotype of this network. The Network Tw-B represented 30.49% of all sequences with four shared haplotypes including Pc_Hap 2 (widespread), Pc_Hap 4 (shared by Tainan, Kaohsiung, and Taichung), Pc_Hap 8 (shared by Pingtung, Yunlin, Hualien), Pc_Hap 5 (shared by Tainan, Pingtung, Chiayi, Taipei, Taichung), and five unique haplotypes from Hualien (Pc_Hap 17, 18), Yilan (Pc_Hap 13, 14) and Chiayi (Pc_Hap 10). Here, Pc_Hap 2 assumed to be the founder haplotype (with most number of connections and a star-like appearance around), The Network Tw-C was represented by only 1.62% of all sequences with one shared haplotype Pc_Hap 9 (Kaohsiung, Hsinchu, Changhua), and one unique haplotype Pc_Hap 7 (Tainan). The distribution Network Tw-C was limited and found from the South (Kaohsiung, Tainan), Middle (Changhua), and North part (Hsinchu). Compared to other cities/counties Tainan, Chiayi, Yilan, and Hualien were reported to have a higher number of shared and unique haplotypes. (Table 1, 2).

**Figure 3.**
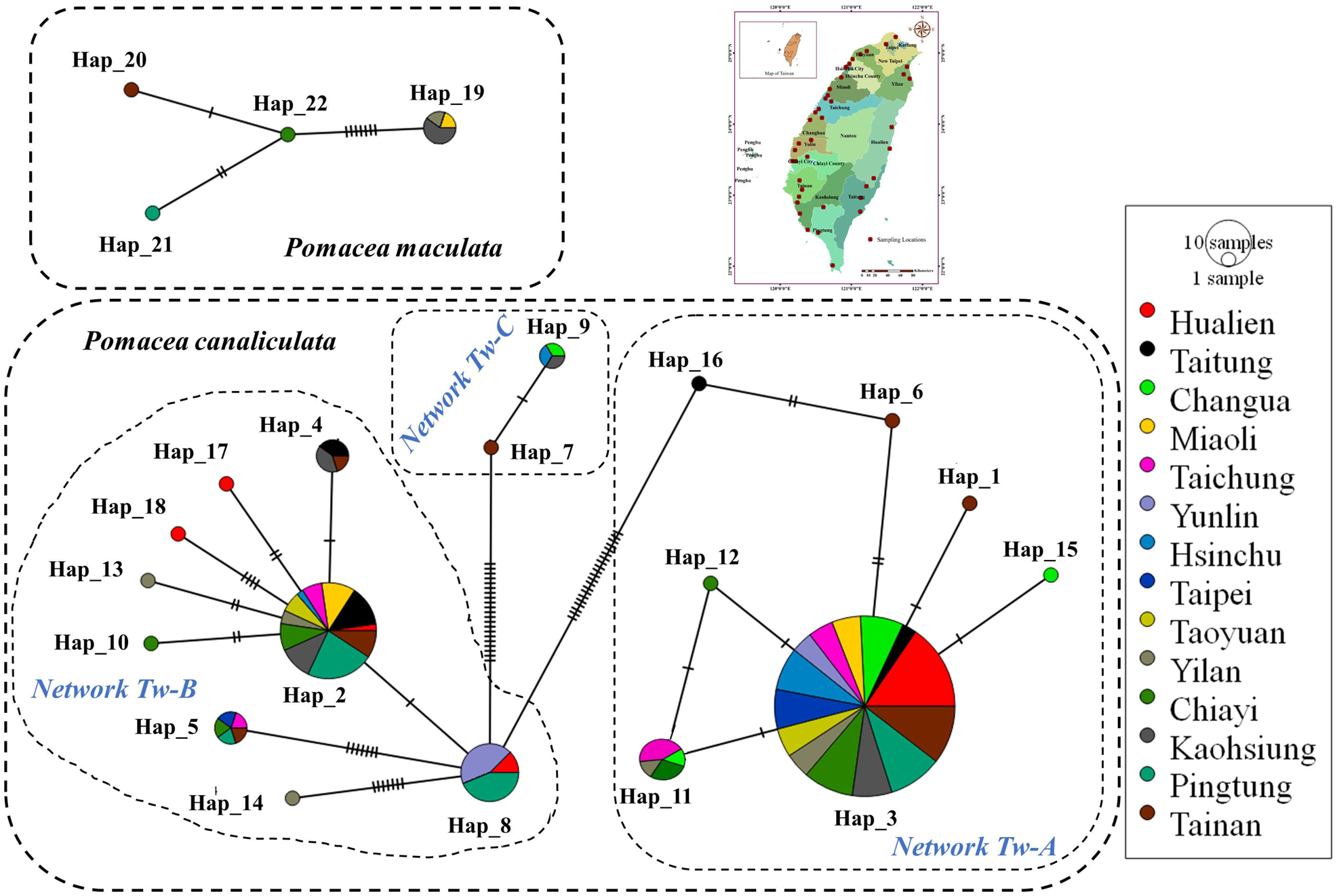
Mitochondrial COI haplotypes networks revealing frequency and relationship of *P. canaliculata* (Pc_Hap 1-18) and *P. maculata* (Pm _Hap 19-22) across Taiwan using Dataset 1.

**Table 1.**
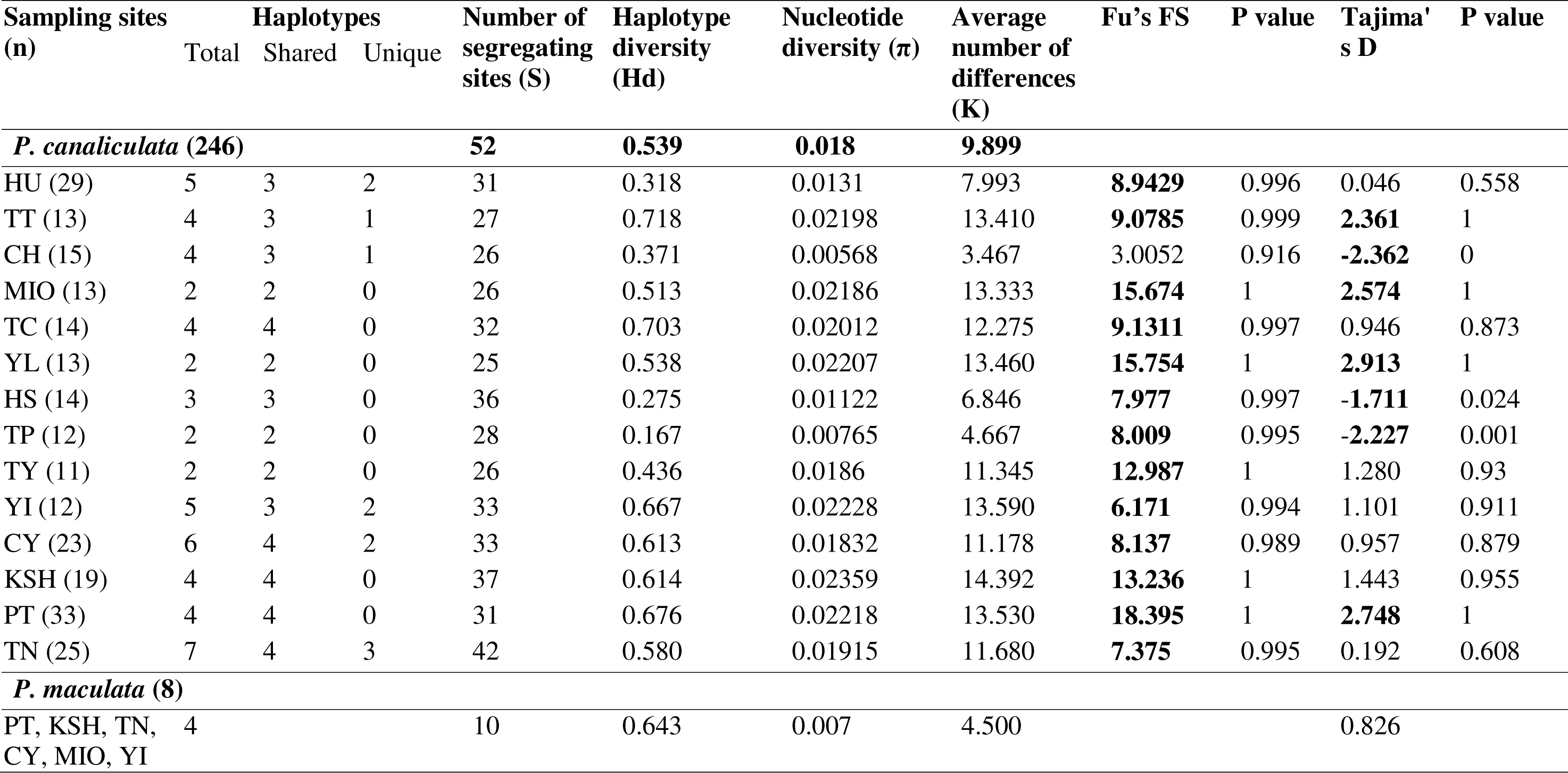
Population genetic diversity of *P. canaliculata* and *P. maculata* across Taiwan. The significant value of Tajima’s *D* and Fu’s *Fs* are marked in bold.

| Sampling sites<br>(n) | Haplotypes | | | Number of<br>segregating<br>sites (S) | Haplotype<br>diversity<br>(Hd) | Nucleotide<br>diversity ($\pi$) | Average<br>number of<br>differences<br>(K) | Fu's FS | P value | Tajima's<br>D | P value |
| --- | --- | --- | --- | --- | --- | --- | --- | --- | --- | --- | --- |
|  | Total | Shared | Unique |  |  |  |  |  |  |  |  |
| ***P. canaliculata* (246)** |  |  |  | **52** | **0.539** | **0.018** | **9.899** |  |  |  |  |
| HU (29) | 5 | 3 | 2 | 31 | 0.318 | 0.0131 | 7.993 | **8.9429** | 0.996 | 0.046 | 0.558 |
| TT (13) | 4 | 3 | 1 | 27 | 0.718 | 0.02198 | 13.410 | **9.0785** | 0.999 | **2.361** | 1 |
| CH (15) | 4 | 3 | 1 | 26 | 0.371 | 0.00568 | 3.467 | 3.0052 | 0.916 | **-2.362** | 0 |
| MIO (13) | 2 | 2 | 0 | 26 | 0.513 | 0.02186 | 13.333 | **15.674** | 1 | **2.574** | 1 |
| TC (14) | 4 | 4 | 0 | 32 | 0.703 | 0.02012 | 12.275 | **9.1311** | 0.997 | 0.946 | 0.873 |
| YL (13) | 2 | 2 | 0 | 25 | 0.538 | 0.02207 | 13.460 | **15.754** | 1 | **2.913** | 1 |
| HS (14) | 3 | 3 | 0 | 36 | 0.275 | 0.01122 | 6.846 | **7.977** | 0.997 | **-1.711** | 0.024 |
| TP (12) | 2 | 2 | 0 | 28 | 0.167 | 0.00765 | 4.667 | **8.009** | 0.995 | **-2.227** | 0.001 |
| TY (11) | 2 | 2 | 0 | 26 | 0.436 | 0.0186 | 11.345 | **12.987** | 1 | 1.280 | 0.93 |
| YI (12) | 5 | 3 | 2 | 33 | 0.667 | 0.02228 | 13.590 | **6.171** | 0.994 | 1.101 | 0.911 |
| CY (23) | 6 | 4 | 2 | 33 | 0.613 | 0.01832 | 11.178 | **8.137** | 0.989 | 0.957 | 0.879 |
| KSH (19) | 4 | 4 | 0 | 37 | 0.614 | 0.02359 | 14.392 | **13.236** | 1 | 1.443 | 0.955 |
| PT (33) | 4 | 4 | 0 | 31 | 0.676 | 0.02218 | 13.530 | **18.395** | 1 | **2.748** | 1 |
| TN (25) | 7 | 4 | 3 | 42 | 0.580 | 0.01915 | 11.680 | **7.375** | 0.995 | 0.192 | 0.608 |
| ***P. maculata* (8)** |  |  |  |  |  |  |  |  |  |  |  |
| PT, KSH, TN,<br>CY, MIO, YI | 4 |  |  | 10 | 0.643 | 0.007 | 4.500 |  |  | 0.826 |  |

**Table 2.**
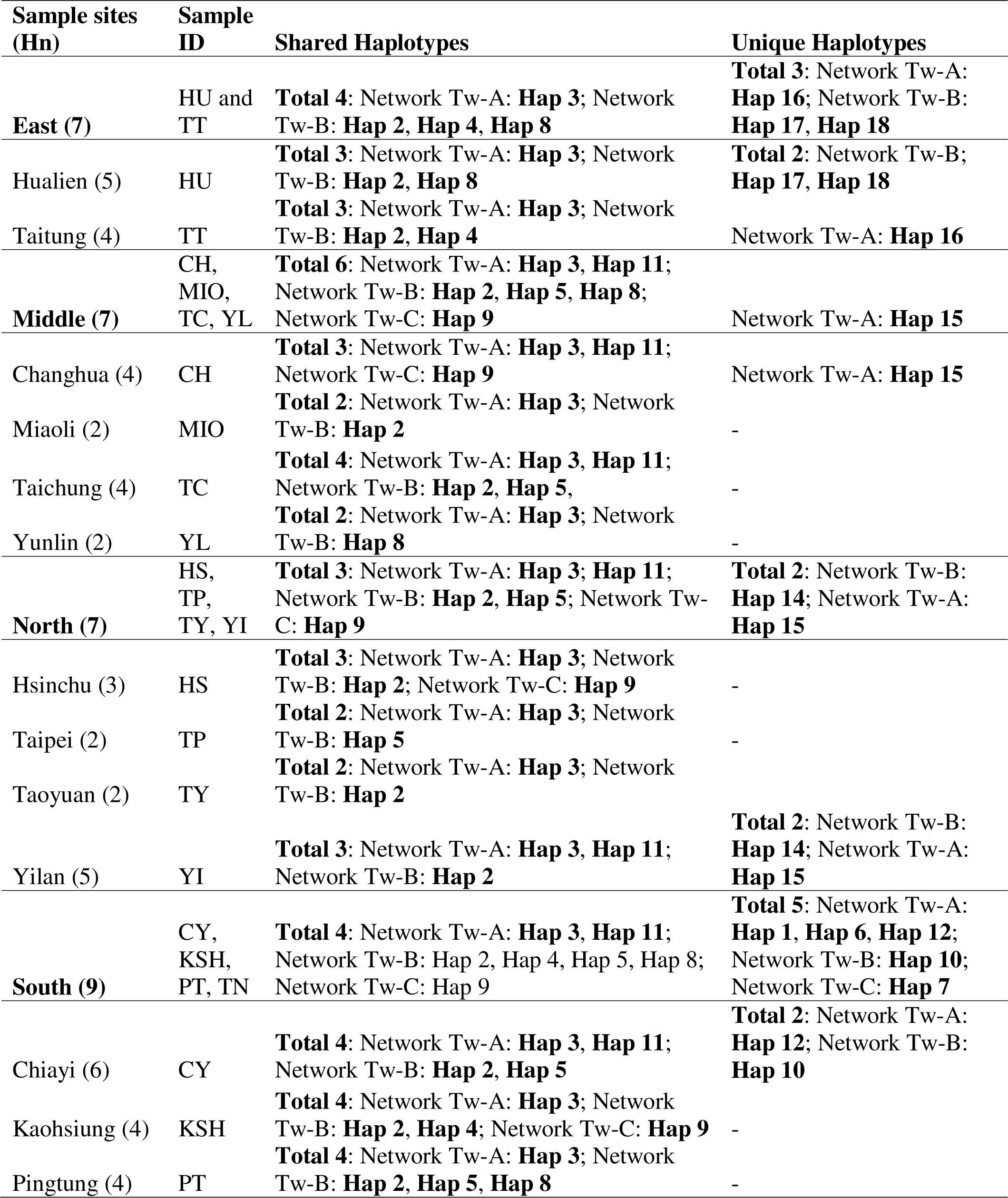
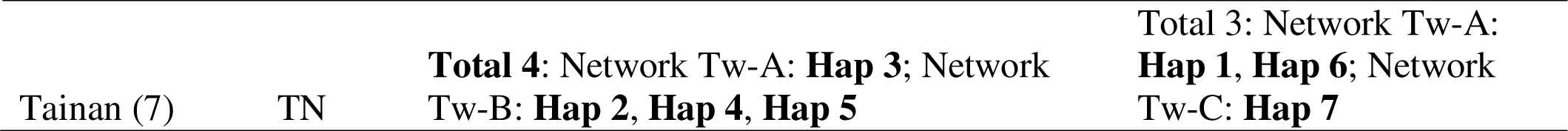
Haplotype distribution (Pc_Hap) of *Pomacea canaliculata* in 14 counties/cities across Taiwan.

#### 3.3.2 Phylogeography of *P. canaliculata* worldwide

The combined mitochondrial COI datasets of *P. canaliculata* with 948 sequences across Africa, Asia, North America, Oceania, and South America were compared to one another and revealed the presence of a total of 65 haplotypes (Wc_Hap1-65) and three distinct networks (Network Wc-A-C) (Fig. 4, APPENDIX 6). The Network Wc-A, representing 32 haplotypes, including the most widely shared dominant Wc_Hap 1, and 31 unique haplotypes from Argentina (Wc_Hap 3-8, 10, 11, 12, 15, 17), Taiwan (Wc_Hap 24, 26, 30, 31, 34), Philippines (Wc_Hap 52, 53, 55, 57, 59, 61), Indonesia (Wc_Hap 48, 49), Thailand (Wc_Hap 62, 63), USA (Wc_Hap 64), China (Wc_Hap 35), Myanmar (Wc_Hap 51), Chile (Wc_Hap 47), Malaysia (Wc_Hap 50). All of those unique haplotypes were connected with Wc_Hap 1, thus inferred as the founder haplotype. This haplotype was also found to be a widespread and dominant haplotype in Taiwan, as well as shared by other Asian countries.

**Figure 4.**
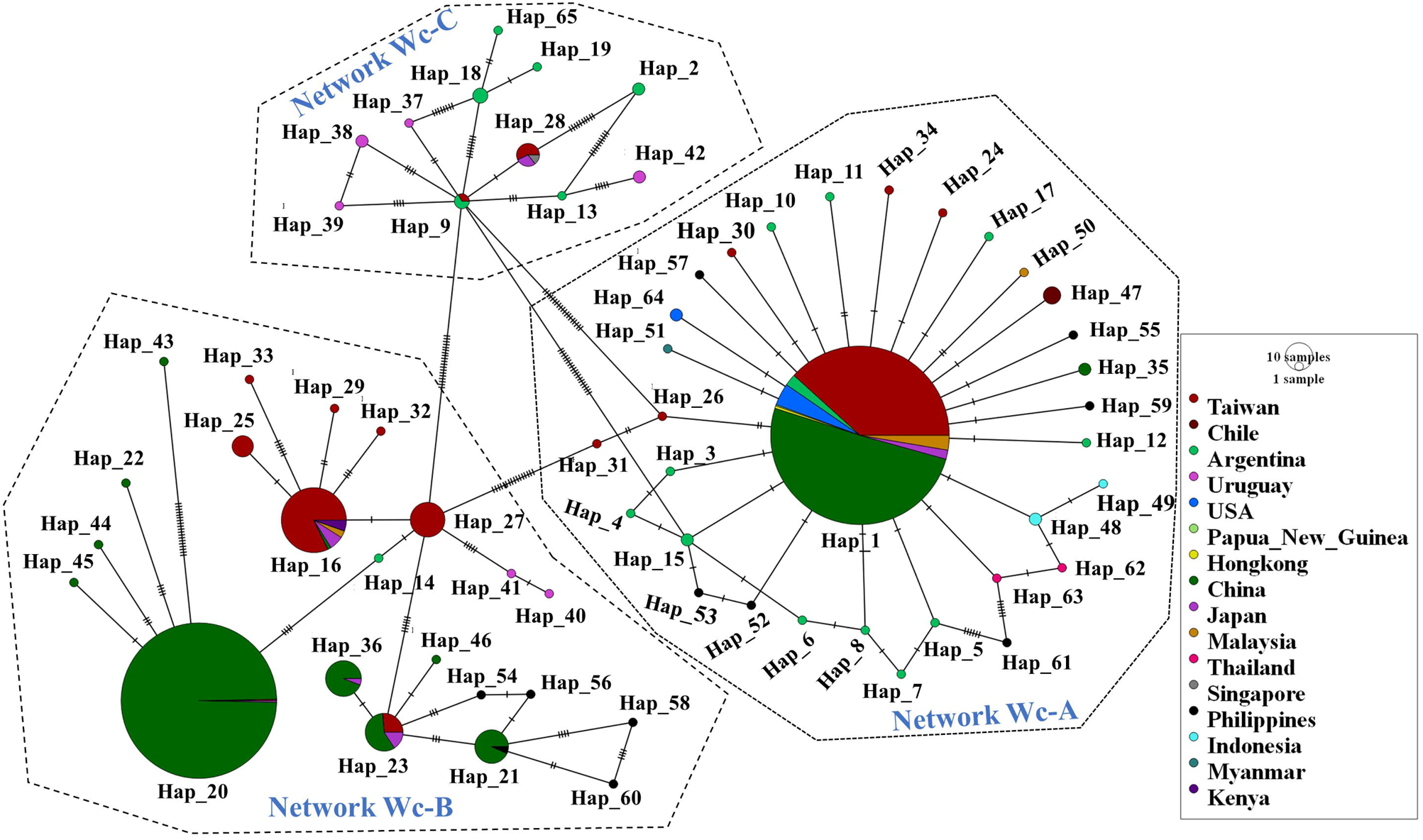
Mitochondrial COI haplotypes (Wc_Hap) revealing frequency and relationship of *P. canaliculata* from the global dataset (Dataset 3).

Network Wc-B represented 22 haplotypes, including five shared; Wc_Hap 16 (Taiwan, China, Japan, Malaysia, and Kenya), Wc_Hap 20 (China, Taiwan, and Philippines), Wc_Hap 21 (China, Philippines), Wc_Hap 23 (Taiwan, China, Japan, Philippines), and Wc_Hap 36 (China and Japan) and 17 unique haplotypes from Taiwan (Wc_Hap 25, 27, 29, 32, 33), China (Wc_Hap 22, 43-46), Uruguay (Wc_Hap 40, 41), Philippines (Wc_Hap 54, 56, 58, 60), and Argentina (Wc_Hap 14). All the shared haplotypes in Network Wc-B were reported from Taiwan, except Wc_Hap 21, and Wc_Hap 36, which were one to three mutational steps away from the shared one. The Wc_Hap 27 (from Taiwan) was noted to be the common connection among all the haplotypes in Network Wc-B.

Lastly, Network Wc-C represented 11 haplotypes, including 2 shared: Wc_Hap 28 (Taiwan, Japan, and Singapore), Wc_Hap 9 (Taiwan and Argentina), and 9 unique haplotypes from Argentina (Wc_Hap 2, 13, 17, 18, 65), and Uruguay (Wc_Hap 37-39, 42). Among the haplotypes in Network Wc-C, Wc_Hap 9 may be inferred as the common ancestor. Most of the haplotypes from introduced regions (Asian countries and African countries) were found to be associated with Network Wc-A and B, however, few representatives were noted in Network Wc-C.

The comparison of representative sequence in Dataset 1 (representing only Taiwan sequences) and Dataset 2 (sequence from Taiwan as well as other native and non-native countries), revealed that haplotypes of Network Tw-A, Network Tw-B, and Network Tw-C, represent in Network Wc-A, Network Wc-B, and Network Wc-C, respectively without any interchange (APPENDIX 6). Taiwan was represented with 18 haplotypes in Dataset 1 (619 bp), however in the global dataset (Dataset 3 = 569 bp; 16 haplotypes from Taiwan) a slight decrease in the number of haplotypes was noted (APPENDIX 7). Across all countries with sequences, the total number of unique haplotypes were found to be the most prevalent in Argentina, Uruguay, Philippines, China, and Taiwan.

#### 3.3.3 Phylogeography of *P. maculata* in Taiwan

In Taiwan, eight sequences of *P. maculata* represented four different haplotypes, where Pm_Hap 19 (62.50%) was the most common and the other three haplotypes (Pm_Hap 20-22) were represented by single sequences (Table 3; APPENDIX 5). Haplotype network analysis of *P. maculata* showed the presence of a single network, where Pm_Hap 1 was a few mutational steps away; however, Pm_Hap 20-22 were more closely related (Fig. 3).

**Table 3.**
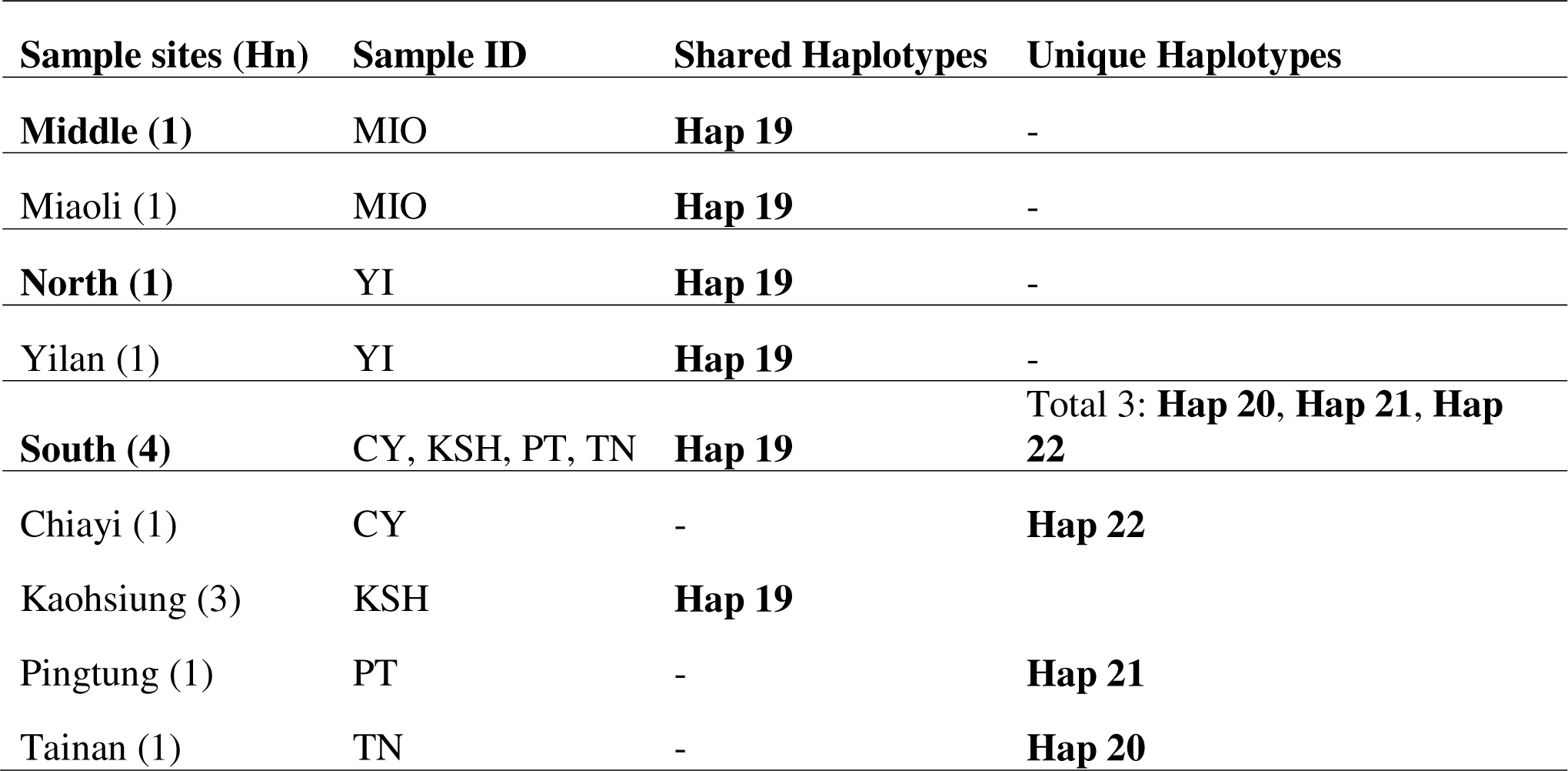
Haplotype distribution (Pm_Hap) of *Pomacea maculata* in 14 counties/cities across Taiwan.

#### 3.3.4 Phylogeography of *P. maculata* worldwide

A total of 226 sequences of worldwide *P. maculata* revealed 46 haplotypes and seven different networks (Network Wm-A-G). Among the haplotype networks, Network Wm-A was widespread, consisting of 29 haplotypes which include four shared haplotypes; Wm_Hap 6 was the most widely distributed (representing 37.61% of all sequences; APPENDIX 8), and Wm_Hap 10 (9.73%) was shared by Argentina and USA, Wm_Hap 9 was shared by USA, Japan, and Argentina, and Wm_Hap 1 was shared by Taiwan, Argentina, and Uruguay, including unique haplotypes from Argentina (Wm_Hap 13-18), Uruguay (Wm_Hap 2-4), Taiwan (Wm_Hap 7, 8), Vietnam (Wm_Hap 5), USA (Wm_Hap 11), and 11 unique to Brazil (Fig. 5). All the other Networks were unique to Brazil. Among the shared haplotypes in Network Wm-A, all other Asian-originated haplotypes created a star-like structure around Wm_Hap 6 and Wm_Hap 1 which suggests those two haplotypes may have been founders throughout Asia. Wm_Hap 6 and Wm_Hap 1 were reported from Taiwan.

**Figure 5.**
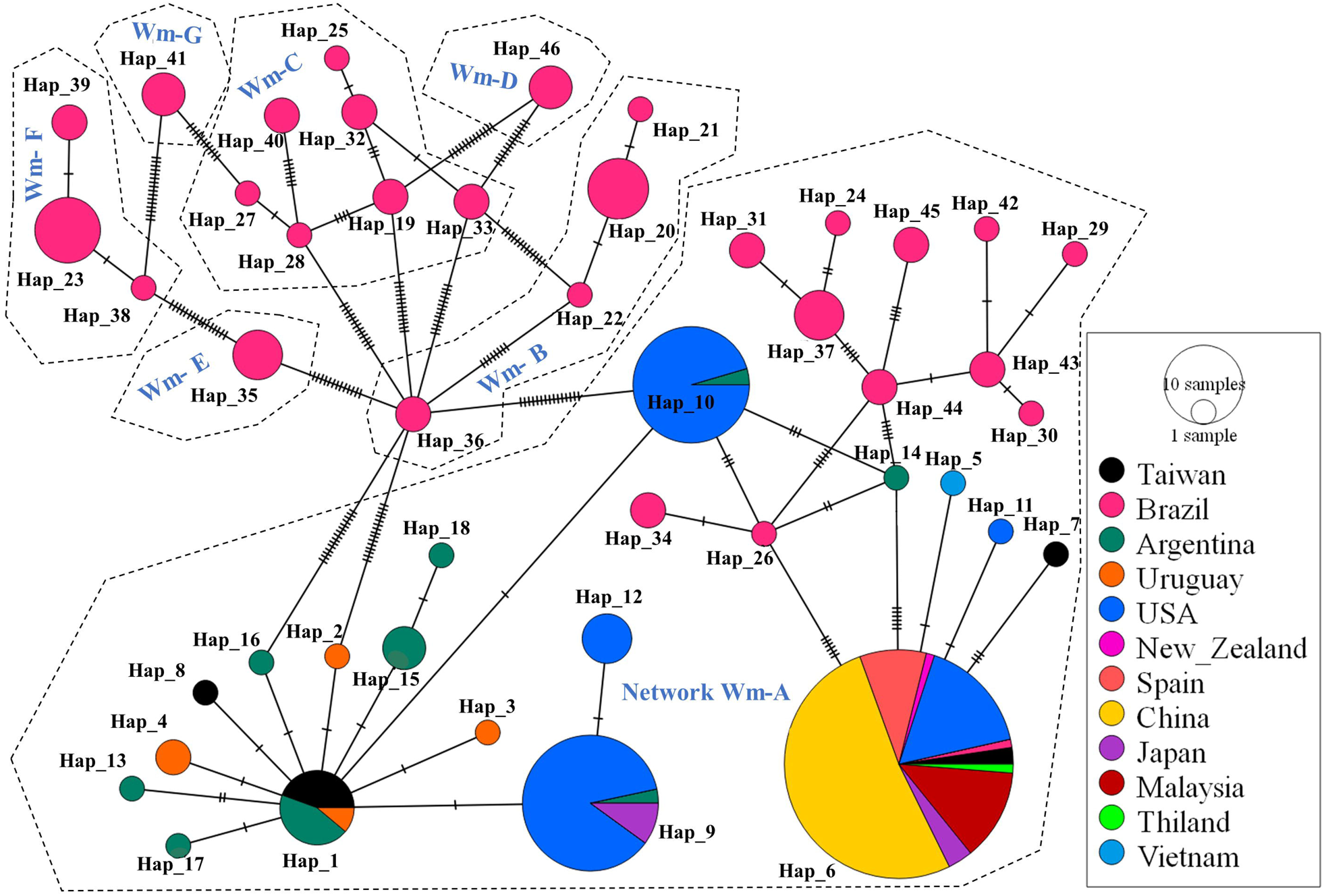
Mitochondrial COI haplotypes (Wm_Hap) revealing frequency and relationship of *P. maculata* from the global dataset (Dataset 2).

### 3.4 Population genetic structure *P. canaliculata* and *P. maculata*

The high value of haplotype diversity (*Hd*), nucleotide diversity (π), and an average number of nucleotide differences (*K*) suggested higher genetic diversity of *P. canaliculata* in Taiwan (Table 1). Among all 14 sites, Taitung, Taichung, Pingtung, Yilan, Kaohsiung, and Chiayi showed the highest genetic diversity; Tainan, Yunlin, Miaoli, and Taoyuan showed high to moderate genetic diversity; Changhua, Hsinchu, Hualien, and Taipei were noted to have less diverse populations (Table 1). The significantly positive value of Fu’s *Fs* and Tajima’s *D* (*D* < 0) were recorded for populations Taitung, Miaoli, Yunlin, Taoyuan, Yilan, Kaohsiung, and Pingtung. Conversely, significantly negative values of Fu’s *Fs* and Tajima’s *D* (*D* > 0) were observed for the population in Changhua, Taipei, and Hsinchu (Table 1). Furthermore, *P. canaliculata* in Taiwan was noted to significantly differ among some provinces, where a higher *F_st_* value (*F_st_* > 0.2) resulted in less gene flow (*Nm* < 1) and vice versa (Table 4). However, the negative *F_st_* values likely indicate more variation within the focal populations than between populations. While *P. maculata* was found to be less dominant in distribution compared to *P. canaliculata*, polymorphism in populations were higher (Table 4).

**Table 4.**
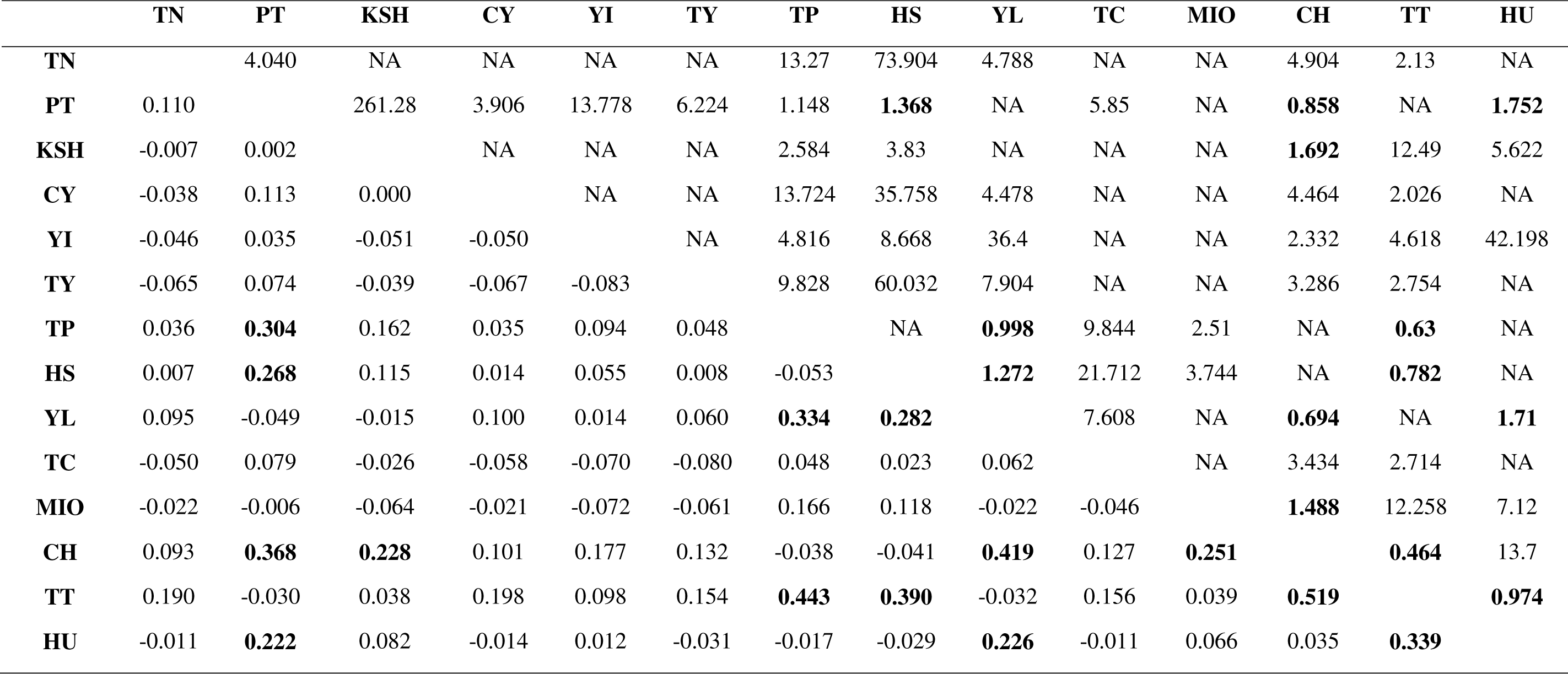
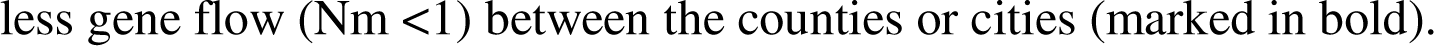
Genetic differentiation coefficient (*F_st_*) below diagonal and gene flow (*N_m_*) above diagonal based on mitochondrial COI sequences of *P. canaliculata* from 14 counties or cities of Taiwan. Abbreviations: HU: Hualien, and TT: Taitung from the East, YL: Yunlin, CH: Changhua, TC: Taichung, and MIO: Miaoli from the Middle, HS: Hsinchu, TY: Taoyuan, TP: Taipei, and YI: Yilan from the North, and PT: Pingtung, KSH: Kaohsiung, TN: Tainan, and CY: Chiayi from the South. The higher *F_st_*value (*F_st_* >0.2) resulted in less gene flow (*Nm* <1) between the counties or cities (marked in bold).

The genetic diversity of *P. canaliculata* and *P. maculata* were considerably higher in their native counties (Argentina, Brazil, and Uruguay) compared to the places in which they were introduced. As an invaded region Taiwan demonstrated a higher genetic diversity of *P. canaliculata* and *P. maculata*, constituting of 16 haplotypes for *P. canaliculata*, and 4 for *P. maculata* (Table 5 and 6).

**Table 5.** Population genetic diversity of *Pomacea canaliculata* from the global mitochondrial COI dataset (Dataset 3).

| Country* | n | Hn | S | Hd | $\pi$ | K |
| --- | --- | --- | --- | --- | --- | --- |
| Argentina | 32 | 19 | 49 | 0.915 | 0.023 | 12.599 |
| Uruguay | 8 | 6 | 35 | 0.928 | 0.027 | 15.107 |
| USA | 19 | 2 | 1 | 0.199 | 0.0003 | 0.199 |
| Chile | 4 | 1 | 0 | 0 | 0 | 0 |
| China | 582 | 12 | 57 | 0.564 | 0.0223 | 12.315 |
| Taiwan | 249 | 16 | 51 | 0.539 | 0.0179 | 9.868 |
| Indonesia | 3 | 2 | 1 | 0.667 | 0.0012 | 0.667 |
| Japan | 18 | 6 | 41 | 0.797 | 0.0277 | 15.274 |
| Malaysia | 14 | 3 | 22 | 0.385 | 0.0105 | 5.780 |
| Philippines | 12 | 12 | 39 | 1 | 0.029 | 16.045 |
| Kenya | 3 | 1 | 0 | 0 | 0 | 0 |
| Thailand | 2 | 2 | 1 | 1 | 0.002 | 1 |
| China-Hongkong** | 2+ 139 | 5 | - | 0.441 | 0.018 | 8.91 |

**Note:** n = Number of sequences; Hn = Number of Haplotype, S = Number of segregating sites; Hd = Haplotype diversity; $\pi$ = Nucleotide diversity; K = Average number of differences
\*Countries represented with 1 sequence were not shown here.
\* 139 samples added from Yang et al., 2022, as their sequence are not available publicly yet.

**Table 6.**
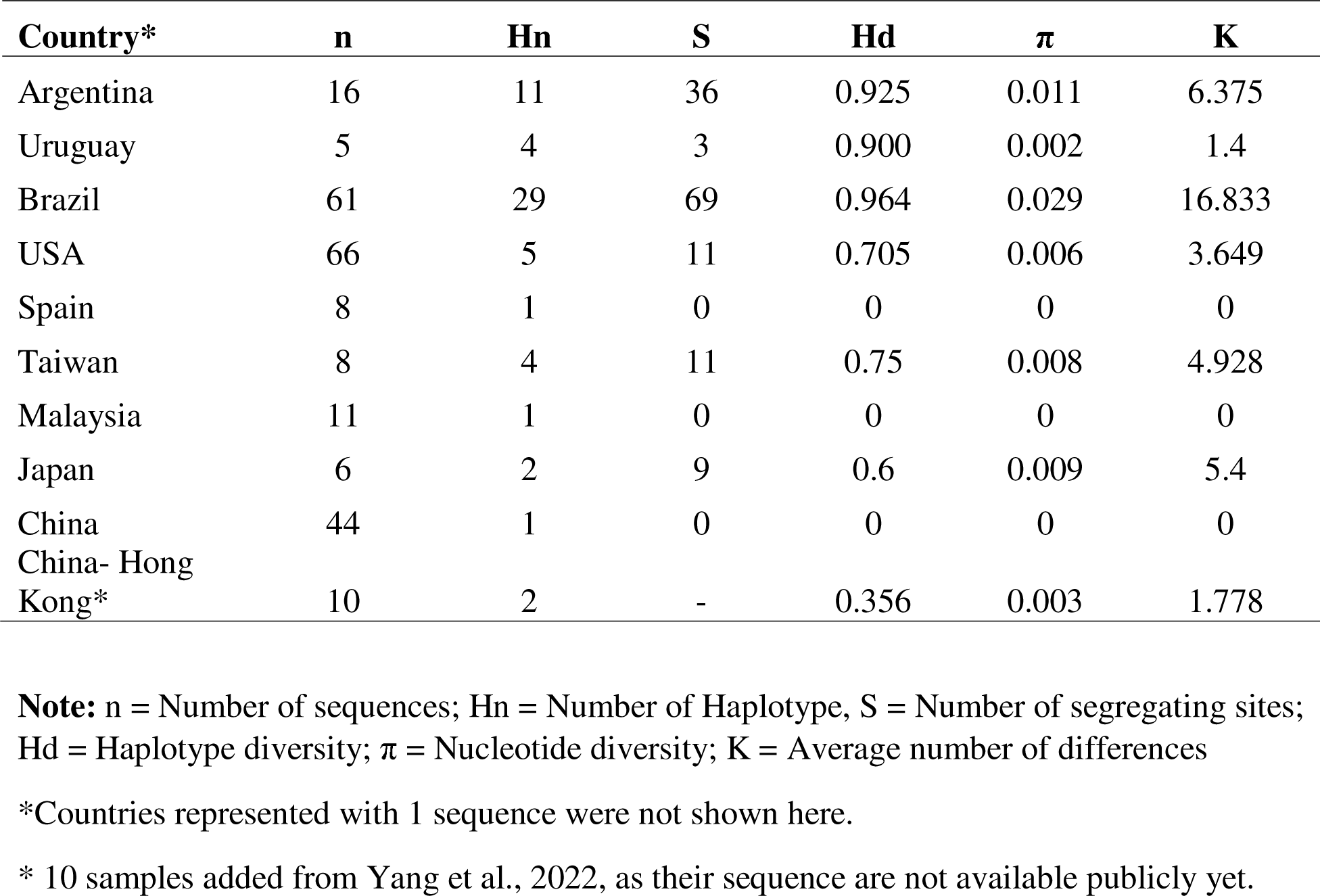
Population genetic diversity of *Pomacea maculata* from the global mitochondrial COI dataset (Dataset 2).

| Country* | n | Hn | S | Hd | $\pi$ | K |
| --- | --- | --- | --- | --- | --- | --- |
| Argentina | 16 | 11 | 36 | 0.925 | 0.011 | 6.375 |
| Uruguay | 5 | 4 | 3 | 0.900 | 0.002 | 1.4 |
| Brazil | 61 | 29 | 69 | 0.964 | 0.029 | 16.833 |
| USA | 66 | 5 | 11 | 0.705 | 0.006 | 3.649 |
| Spain | 8 | 1 | 0 | 0 | 0 | 0 |
| Taiwan | 8 | 4 | 11 | 0.75 | 0.008 | 4.928 |
| Malaysia | 11 | 1 | 0 | 0 | 0 | 0 |
| Japan | 6 | 2 | 9 | 0.6 | 0.009 | 5.4 |
| China | 44 | 1 | 0 | 0 | 0 | 0 |
| China- Hong Kong* | 10 | 2 | - | 0.356 | 0.003 | 1.778 |

**Note:** n = Number of sequences; Hn = Number of Haplotype, S = Number of segregating sites; Hd = Haplotype diversity; $\pi$ = Nucleotide diversity; K = Average number of differences
\*Countries represented with 1 sequence were not shown here.
\* 10 samples added from Yang et al., 2022, as their sequence are not available publicly yet.

### 3.5 Mismatch distribution

The mismatch distribution results of *P. canaliculata* in Taiwan showed two bimodal well-separated peaks with some small intermediate peaks (Fig. 6a). However, in the case of *P. maculata* three distinct peaks were observed, including two major peaks and an intermediate peak (Fig. 6b). The mismatch distribution of *P. canaliculata* and *P. maculata* from the world dataset showed multimodal distribution (Fig. 6c and d).

**Figure 6.**
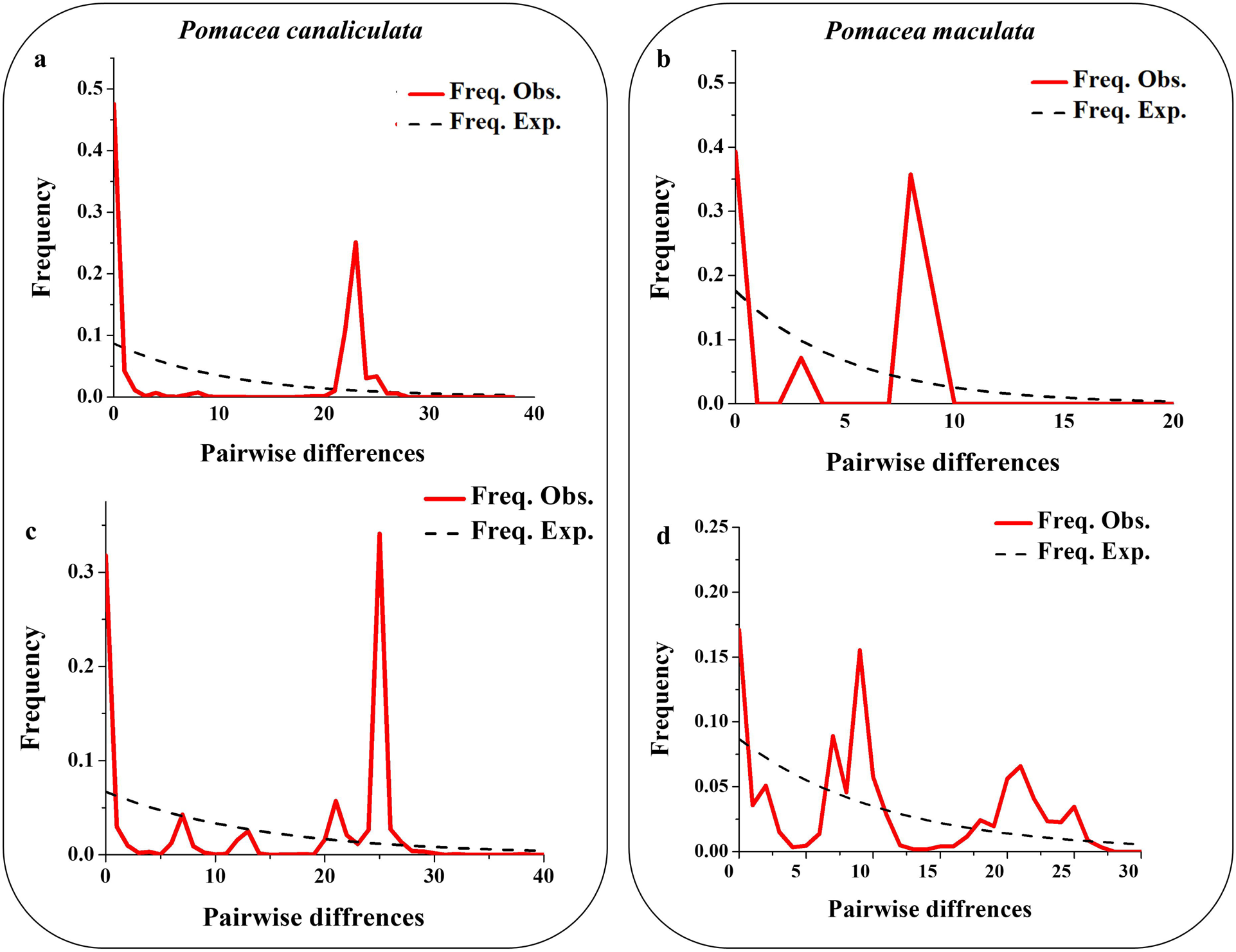
Mismatch distribution analysis of *P. canaliculata* from (a) Taiwan (Dataset 1) and (c) the global dataset (Dataset 3), and *P. maculata* (b) Taiwan (Dataset 1) and (d) the global dataset (Dataset 2)

## 4. DISCUSSION

Based on our mitochondrial COI barcoding data, we found a wide distribution of *P. canaliculata* across Taiwan, whereas *P. maculata* was less abundant and restricted few counties (Pingtung, Kaohsiung, Tainan, and Chiayi, Miaoli and Yilan) (Fig. 1). Moreover, the distribution of *P. maculata* and *P. canaliculata* were noted to be sympatric and heterogenous across the country (Fig. 1 and Table 1), a similar distribution pattern was noted in our previous study in southern Taiwan (Banerjee et al., 2022). Our haplotype analysis further demonstrated the possible origin of *P. maculata* from Argentina, although *P. maculata* may have been introduced independently from Argentina and Brazil (Hayes et al., 2008; Yang et al., 2022) instead of being imported directly from Brazil as reported in cases of *P. maculata* populations in China (Yang et al., 2018). Thus, the results from Yang et al. (2018, 2022) and this present study showed that the origin of *P. maculata* may be different in mainland China from Hong Kong and Taiwan.

Interestingly, the invasion of *P. canaliculata* and *P. maculata* has also been noted to be sympatric and heterogenous in other nearby Asian countries, such as in China (Yang et al., 2018), Hong Kong (Yang et al., 2022), Malaysia (Rama Rao et al., 2018), and Thailand (Dumidae et al., 2021), where *P. canaliculata* is also reported to be more widely distributed compared to *P. maculata*. This corroborates the dominant hypothesis of predominately *P. canaliculata* in Asia regions (Hayes et al., 2008). Interestingly, this distribution pattern is not reflected in their native ranges, where *P. maculata* is routinely found to be widely distributed throughout Brazil, Uruguay, and Argentina, whereas *P. canaliculata* remains restricted to North Argentina and South Uruguay (Hayes et al., 2008; Glasheen et al., 2020). The difference in the distribution in Asian regions may reflect different physiological tolerance, or species-specific biology (Hayes et al., 2008; Matsukura et al, 2016). Alternatively, it can be posited that these phylogeographical patterns and unequal distribution may be due to complex introduction histories. Notably, current distribution data is based on mitochondrial DNA, and *P. canaliculata* and *P. maculata* are reported to hybridized both in their native and non-native ranges (Glasheen et al., 2020; Yang et al., 2020; Kannan et al., 2021). Thus, hybridization could also play an important role in their unequal phylogeographic distribution. Similar to *P. maculata*, another invasive species, *P. scalaris,* was previously reported to be restricted to the southern part of Taiwan (Wu et al., 2010), however, we recorded some individuals from the middle region of Taiwan during our collection, possibly suggesting slow expansion.

Although the origin of *P. canaliculata* and *P. maculata* in Asia has been widely studied by previous researchers from different countries (Hayes et al., 2008; Liu et al., 2019; Dumidae et al., 2021; Yang et al., 2022), additional data from Taiwan specifically (and for the first time) will contribute to our knowledge regarding the possible path of introduction and spread. Our haplotype analysis revealed the presence of all three independent networks of *P. canaliculata* across Taiwan, whereas only two networks of *P. canaliculata* were detected from other countries of Asia (Yang et al., 2018; Yang et al., 2021). The occurrence of three largely different networks across Taiwan suggesting introduction may have occurred multiple times and possibly propagated from multiple locations (Fig. 3).

The combined mitochondrial datasets of *P. canaliculata* with 948 global sequences (originating from Africa, Asia, North America, Oceania, and South America) exhibited three different networks, mirroring previous studies (Yang et al., 2018; Yang et al., 2021), with sequences from Taiwan represented across all networks (Fig. 4). The Wc_Hap 1, Wc_Hap 27, and Wc_Hap 9 represented the founder haplotypes of Network Wc-A, Network Wc-B, and Network Wc-C respectively. The common sharing of founder haplotypes between Taiwan and Argentina reaffirms the historical assumption of the introduction of *P. canaliculata* from Argentina to Taiwan (Joshi & Sebastian, 2003; Hayes et al., 2008). Although the absence of a shared haplotype from Argentina was noted in the case of WC_Hap 27 in Network Wc-B, which again may simply be due to a smaller number of represented sequences from Argentina in our global dataset. Furthermore, Network Wc-A and Network Wc-B were widely distributed, however the haplotypes from Network Wc-C were rare in all native and non-native countries, not just those haplotypes representative in Asia. This also does not lend support for sequence sampling bias (i.e. more representative samples in Taiwan compared to other sampled countries), although we cannot preclude it as a possible contributing factor.

Interestingly, our haplotype analysis revealed the presence of a single Network for *P. maculata* across Taiwan, echoing patterns across other Asian countries (Yang et al., 2018; Yang et al., 2021). This is in contrast to *P. canaliculata* for which the haplotype distribution pattern suggests multiple introduction events (Fig. 3 and 5). Corroborating this further and despite the complex network of unique haplotypes, Taiwan shares the founder haplotype with Argentina (Wm_Hap 1). Notably, our results demonstrated the presence of Brazilian haplotypes with only a few mutational steps away from the founder haplotype (Wm_Hap 1), suggesting the introduction of *P. maculata* possibly originated via Argentina and Brazil, rather than solely from Brazil (Fig. 5). Similar findings were also noted in Hong Kong, where results from Yang et al. (2022) demonstrated *P. maculata* was possibly introduced from Argentina and Brazil. The presence of only a single network of *P. maculata* may suggest less frequent introductions events in Taiwan.

Unsurprisingly, the populations of *P. canaliculata* and *P. maculata* in Taiwan showed lower haplotype diversity compared to their native population counterparts (e.g., Argentina and Brazil) which can be the result of founder effects. However, compared with other Asian countries, Taiwan showed similar (e.g., China) or higher (e.g., Malaysia, Thailand, and Philippines) genetic diversity for *P. canaliculata* and *P. maculata* (Table 5 and 6). These observations insinuate that Taiwan may have faced multiple introductions during the past. Furthermore, as per our mismatch distribution analysis, two distinct peaks with large pairwise differences created a well-separated bimodal mismatch graph for *P. canaliculata* (Fig. 6a), however mismatch distribution analysis produced an intermediate peak in between the bimodal graph for *P. maculata* (Fig. 6b). Those patterns suggest two episodes of population expansion which may coincide with multiple introductions for *P. canaliculata*, and possibly a single or continuous source for *P. maculata*. Furthermore, the multimodal mismatch distribution of *P. canaliculata* and *P. maculata* for the global dataset (Dataset 2 and 3) indicated their complex population expansion dynamics, and possibly introduction history, around several countries (Fig. 6c and d).

Their genetic diversity (value of *Hd*, π, and *K*) varied among the counties/cities of Taiwan which may be related to multiple sources of introductions at different times, across different land-use patterns (e.g., agriculture), and human transportation. Moreover, the dispersal ability of *Pomacea* spp. is observed to be low and thus the widespread distribution across Taiwan within such a short invasion period was probably facilitated predominantly from human transportation.

The neutrality test, such as the value of Fu’s *Fs* and Tajima’s *D* (D < 0), was not observed as significant for the overall population *P. canaliculata* and *P. maculata* in Taiwan, however, the higher value of Fu’s *Fs* and Tajima’s *D* (*D* < 0) for the population of *P. canaliculata* in Taitung, Miaoli, Yunlin, Taoyuan, Yilan, Kaohsiung, and Pingtung did demonstrate a high number of rare alleles. On the contrary, the lower value of Fu’s *Fs* and Tajima’s *D* (*D* > 0) for the population of *P. canaliculata* in Changhua, Taipei, and Hsinchu suggests a scarcity of rare alleles, however, these results may equally be due to sampling error. Furthermore, the significant difference in *F_st_* value (*F_st_*> 0.2) and negative *F_st_* values (suggest more variation within the populations than in between) in Table 4 could be result of multiple introductions, human transportation and population mixing.

Although the mitochondrial DNA haplotyping data is useful to understand the genetic diversity, as *P. canaliculata* and *P. maculata* are reported to hybridize, implementing nuclear genomic data or whole genomic sequencing data across invaded and native countries will be more useful for understanding invasion dynamics. The current datasets were limited in number and range of collections across different countries or regions, which could be solved by adding more precisely identified sample sequences.

## Conclusions

Our data suggested the Argentinian origin of *P. canaliculata* in Taiwan, whereas *P. maculata* might have both Argentinian and Brazilian origin. Comparing their introduction histories, current distribution, and cryptic nature of *P. canaliculata* and *P. maculata*, we assume that *P. maculata* may have been introduced along with *P. canaliculata* (either intentionally or unintentionally) in Taiwan, however, *P. canaliculata* likely has several introduction histories than *P. maculata*. Furthermore, our sampling across Taiwan, also reported the presence of *P. scalaris* (non-target for this study) within the middle region of Taiwan, which possibly suggest the expansion of another invasive species. Thus, the establishment of *Pomacea* spp. across all of Taiwan is alarming, and our current data suggests the urgent need for a nationwide mitigation program and proper management policy. For early detection and to prevent further introductions and spread of multiple *Pomacea* species, eDNA-based methods have been previously demonstrated to be useful (Banerjee et al., 2022).

## ACKNOWLEDGEMENTS

P.B. has been supported by Overseas Research Scholarships (ORS) from National Chung Cheng University as well as Ministry of Education (MOE)-Industry-Academia project (Taiwan). The authors would also like to thank the Ministry of Science and Technology (Taiwan) for financial support (MOST 109-2811-M-194-502; MOST 108-2811-M-194-510).

## AUTHORS’ CONTRIBUTIONS

P.B. and C.Y.C. conceived of the idea; P.B., performed experiments, P.B., K.A.S., and C.Y.C., prepared the first draft and revised the manuscript, figures and tables. G.D., R.K.S, J.P.M., and M.W.Y.C, K. H. L., gave extensive edits and revised the manuscript. We would like to thank Prof. Qian-Qian Yang (College of Life Sciences, China Jiliang University, Hangzhou 310018, China) for her valuable suggestions during revisions.

## DATA AVAILABILITY STATEMENT

All data obtained in this study have been deposited to GeneBank under the accession number OQ977961 - OQ978203 and OQ977020-OQ977027.

---

**APPENDIX 1.**
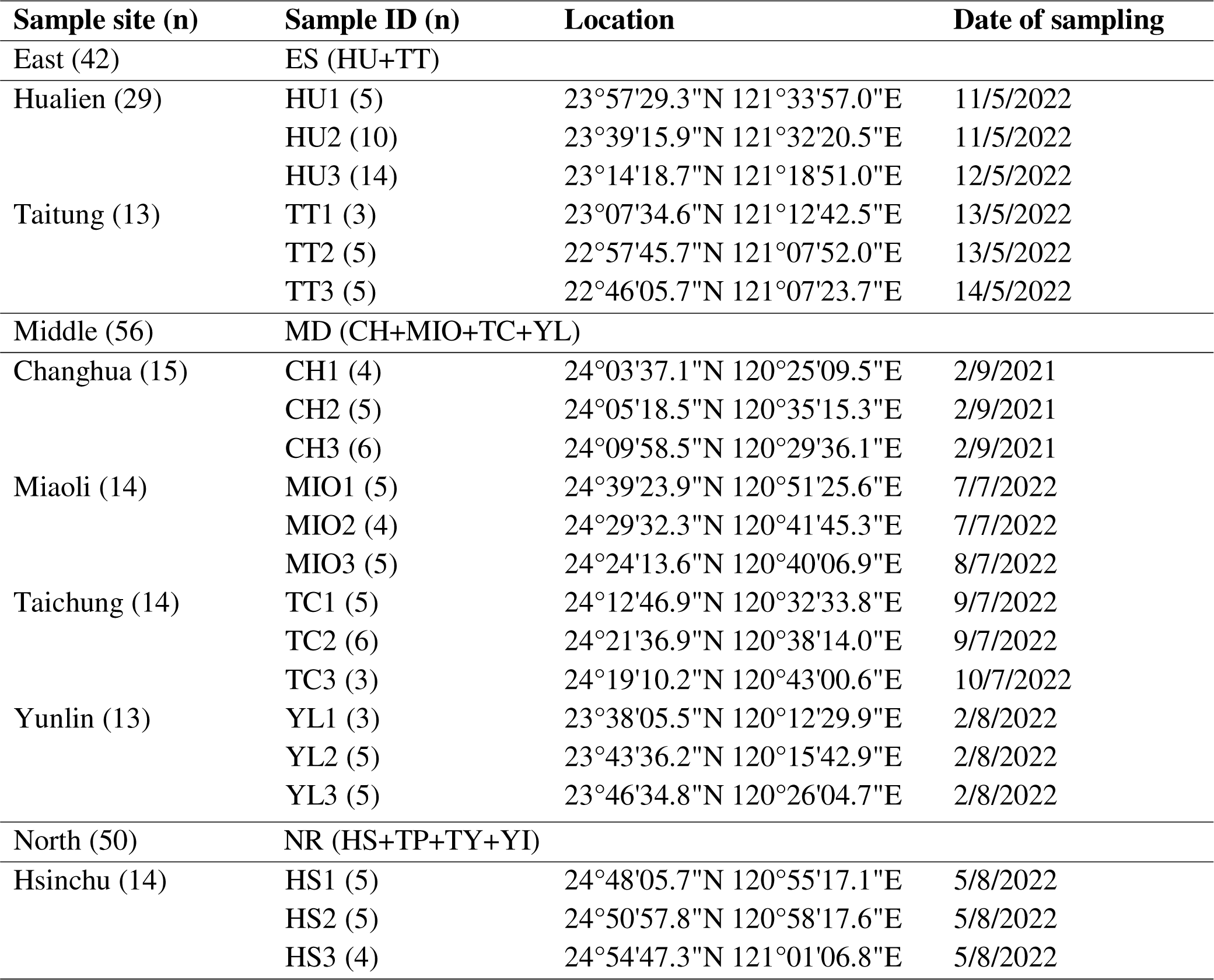
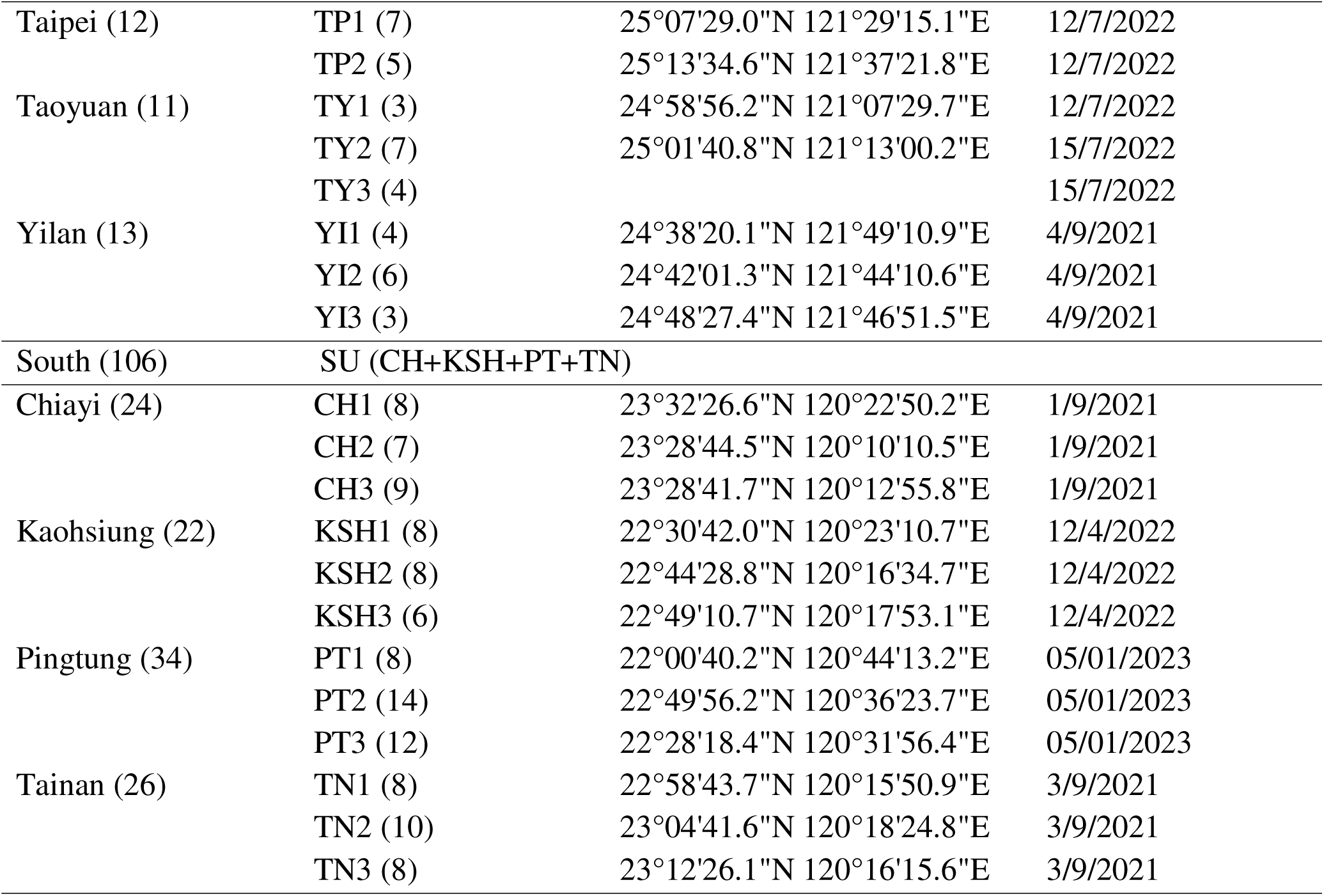
Details of sampling sites, number of samples (n), and sampling locations across Taiwan for *Pomacea* spp.

**APPENDIX 2.**
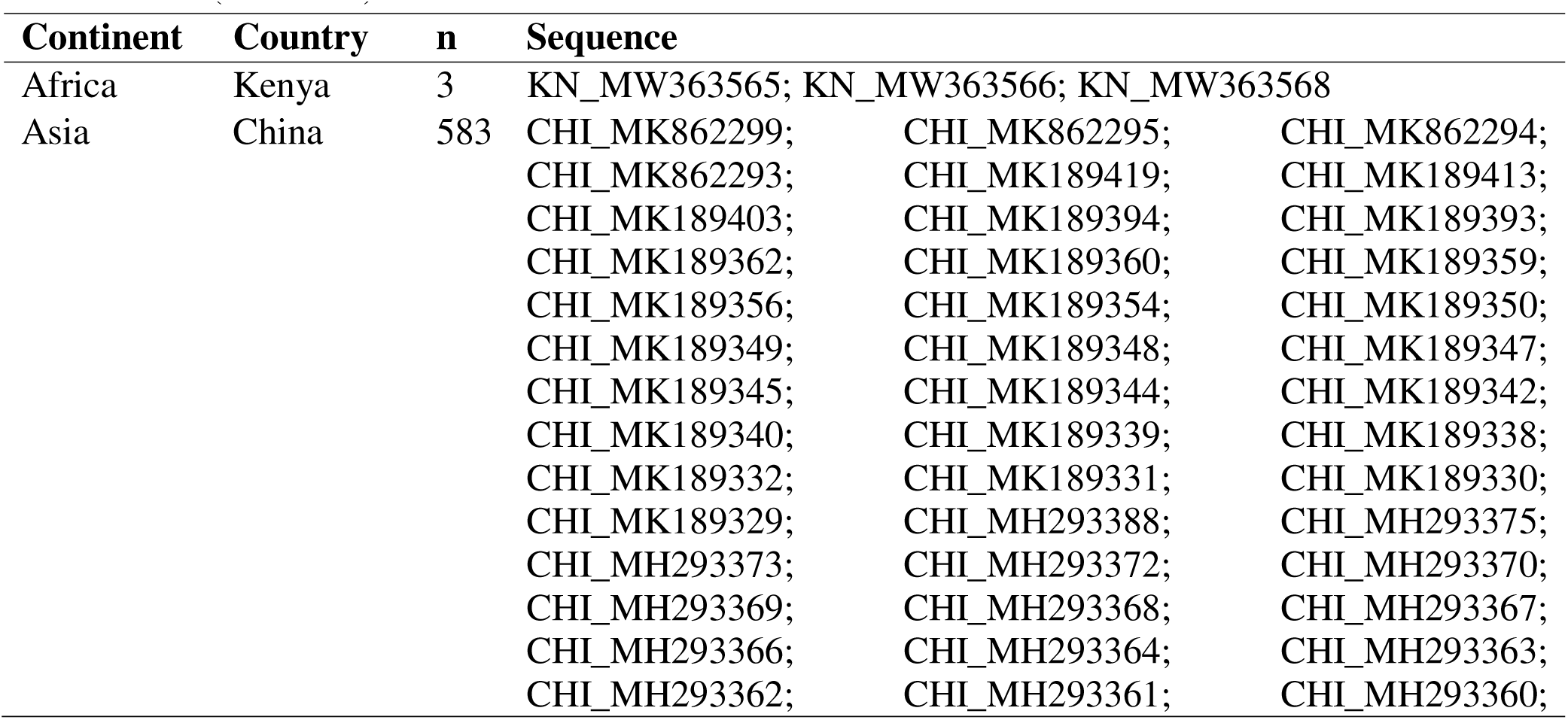
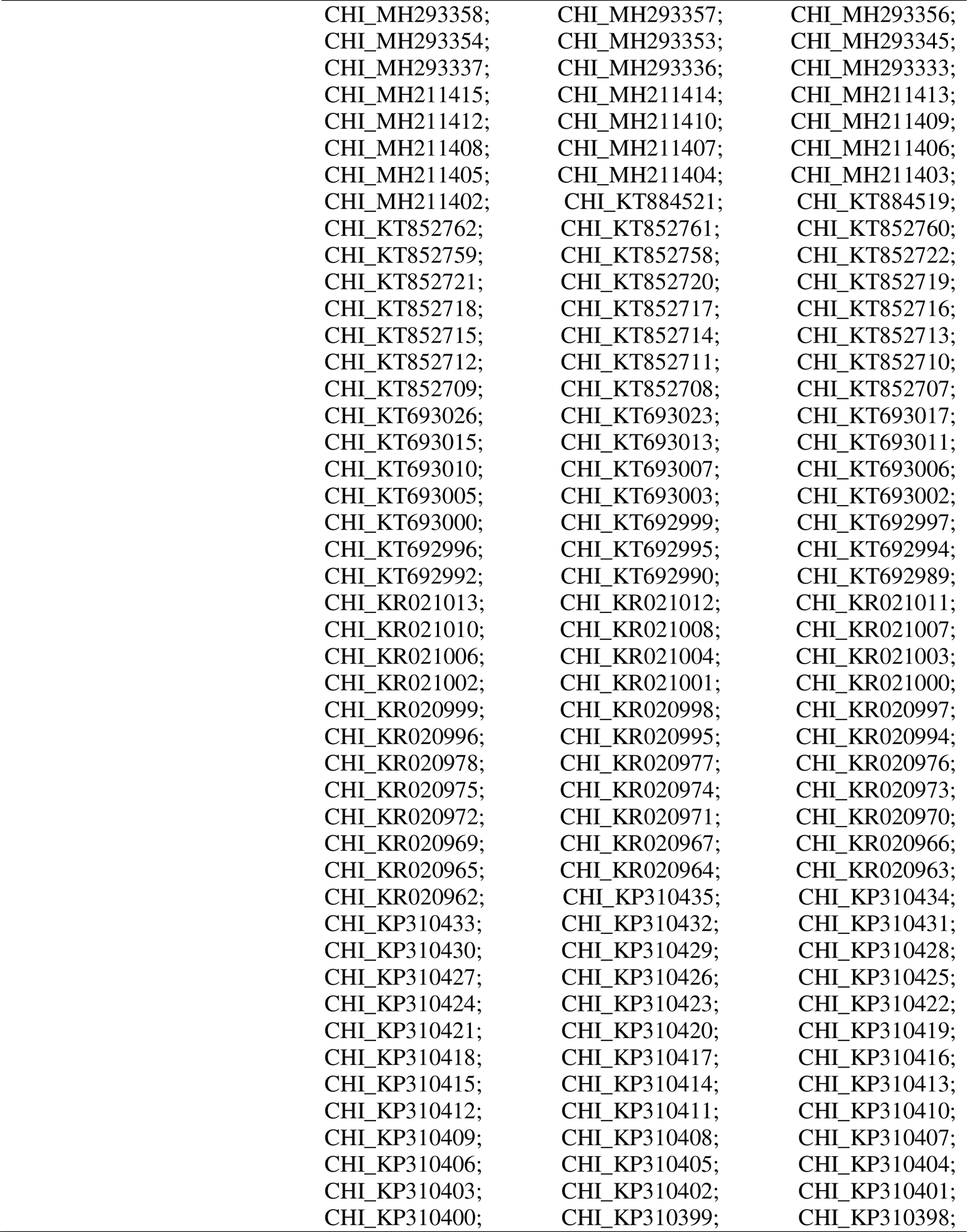
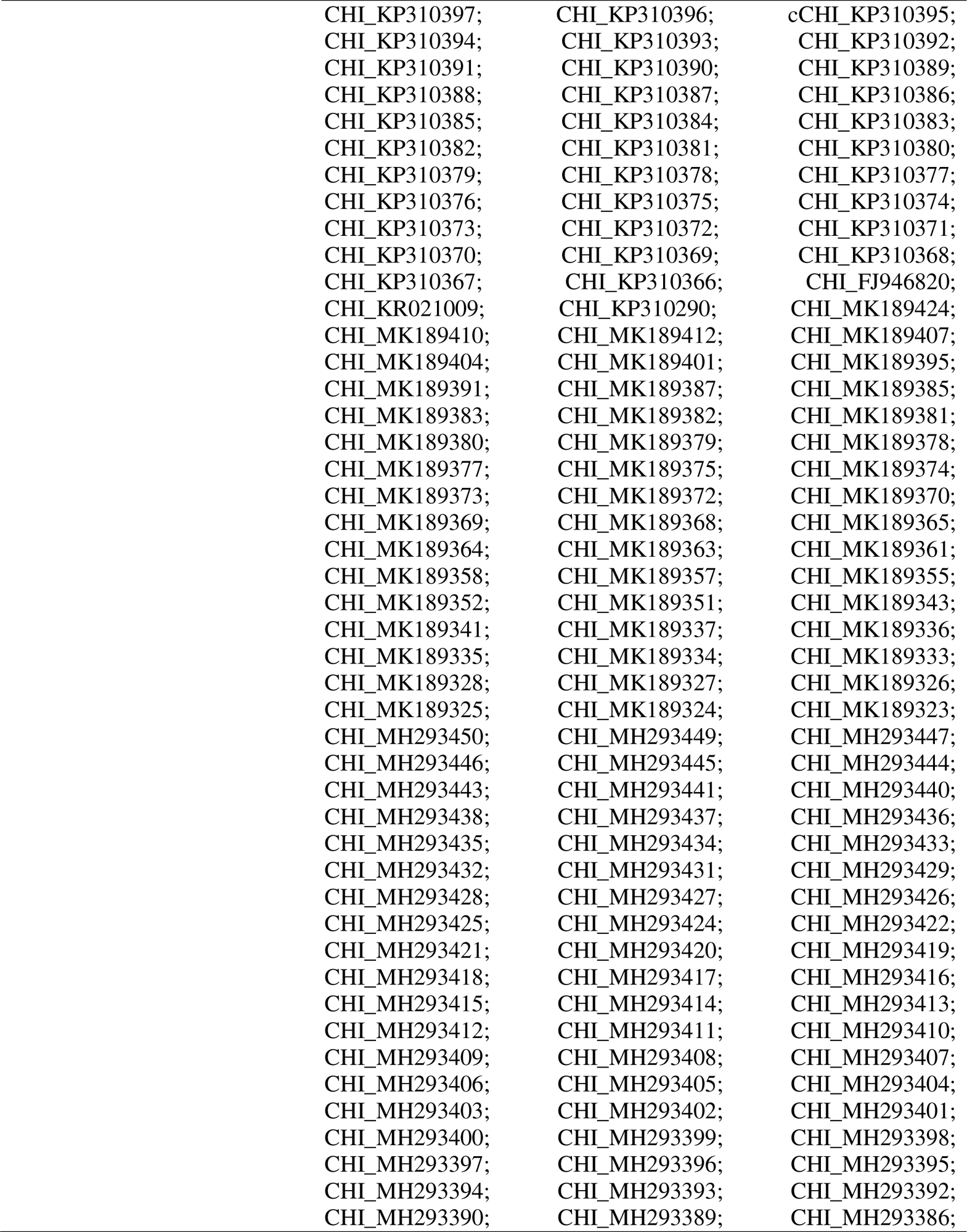
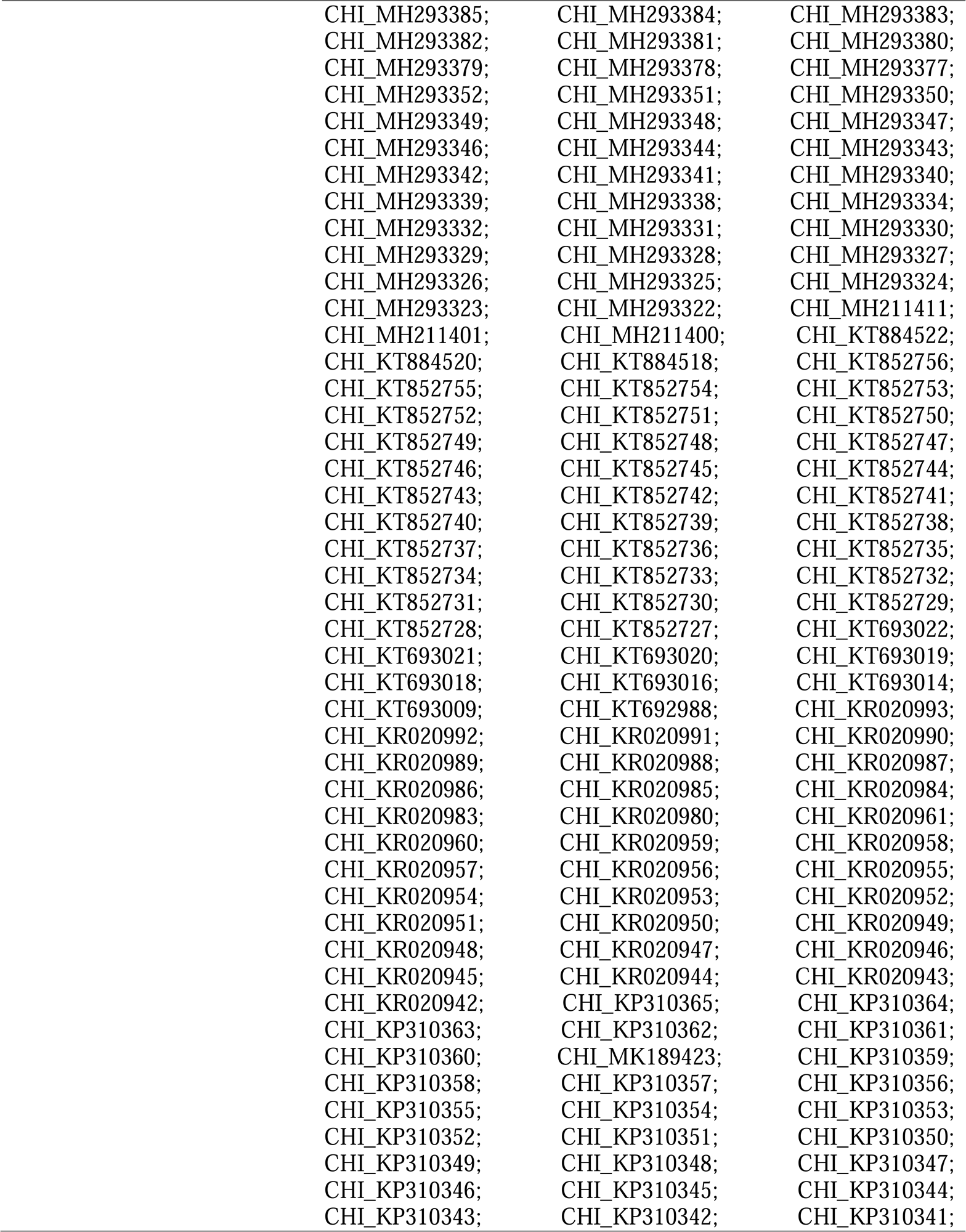
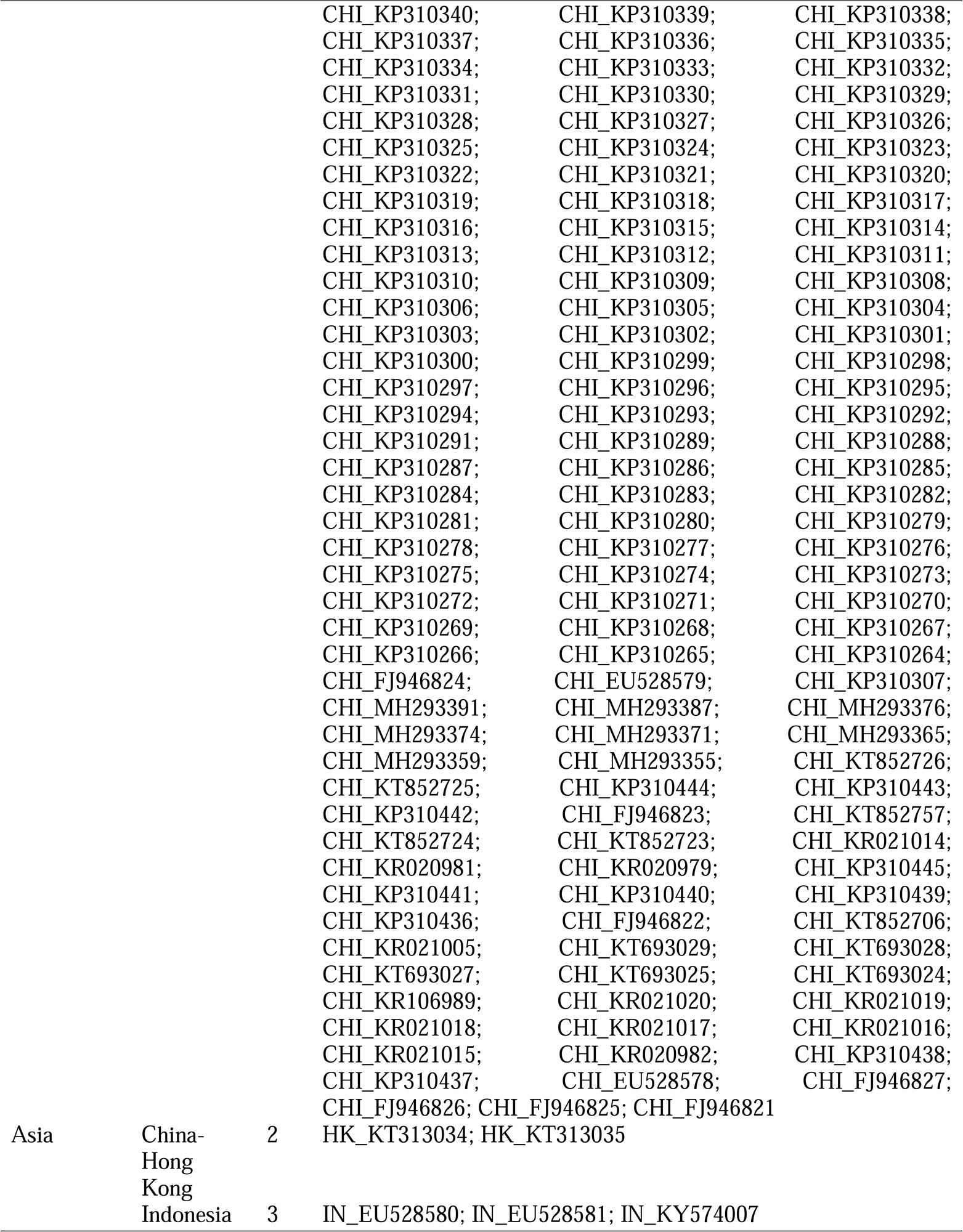
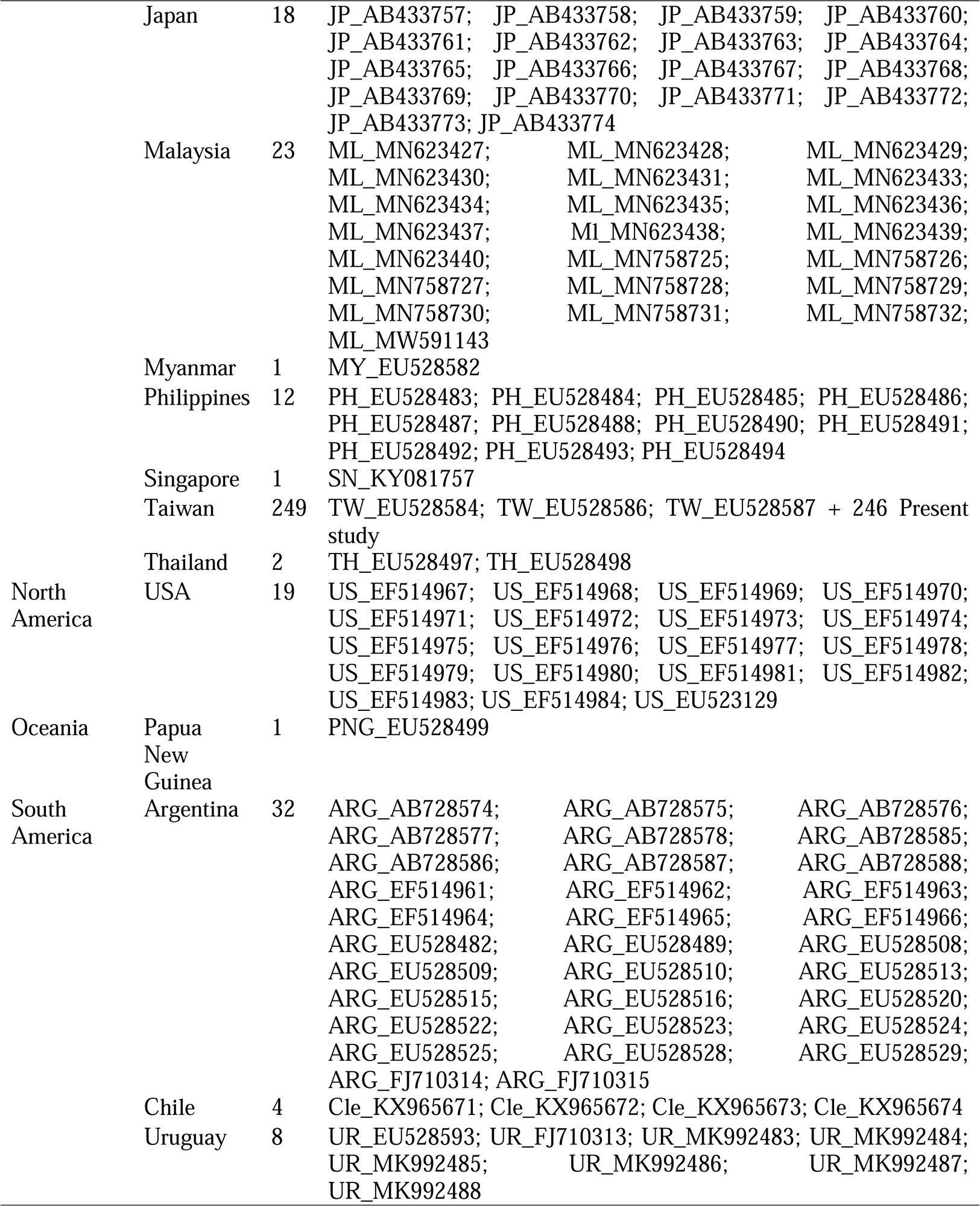
List of GenBank accession numbers used to prepare *Pomacea canaliculata* global mitochondrial COI dataset (Dataset 3)

**APPENDIX 3.**
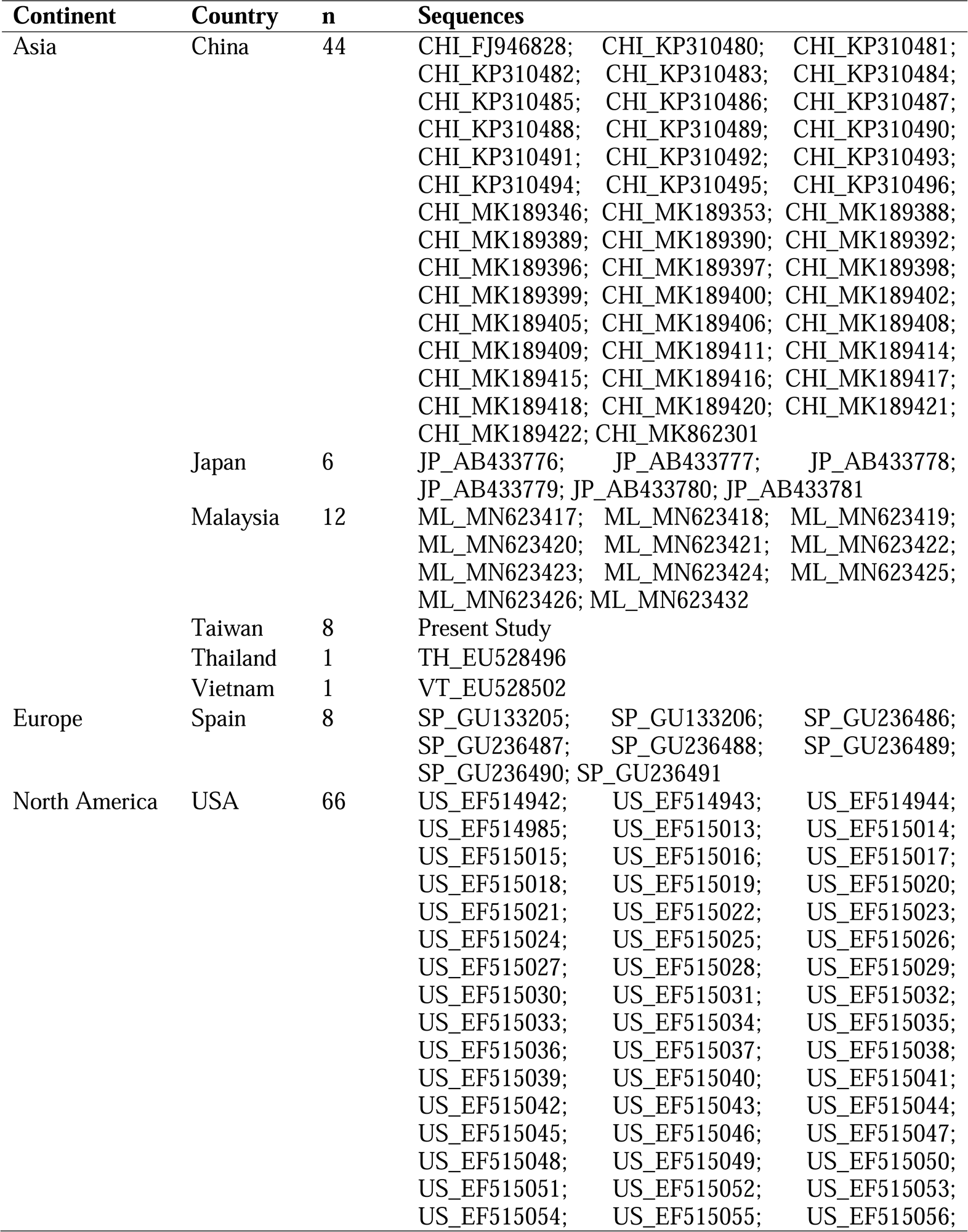
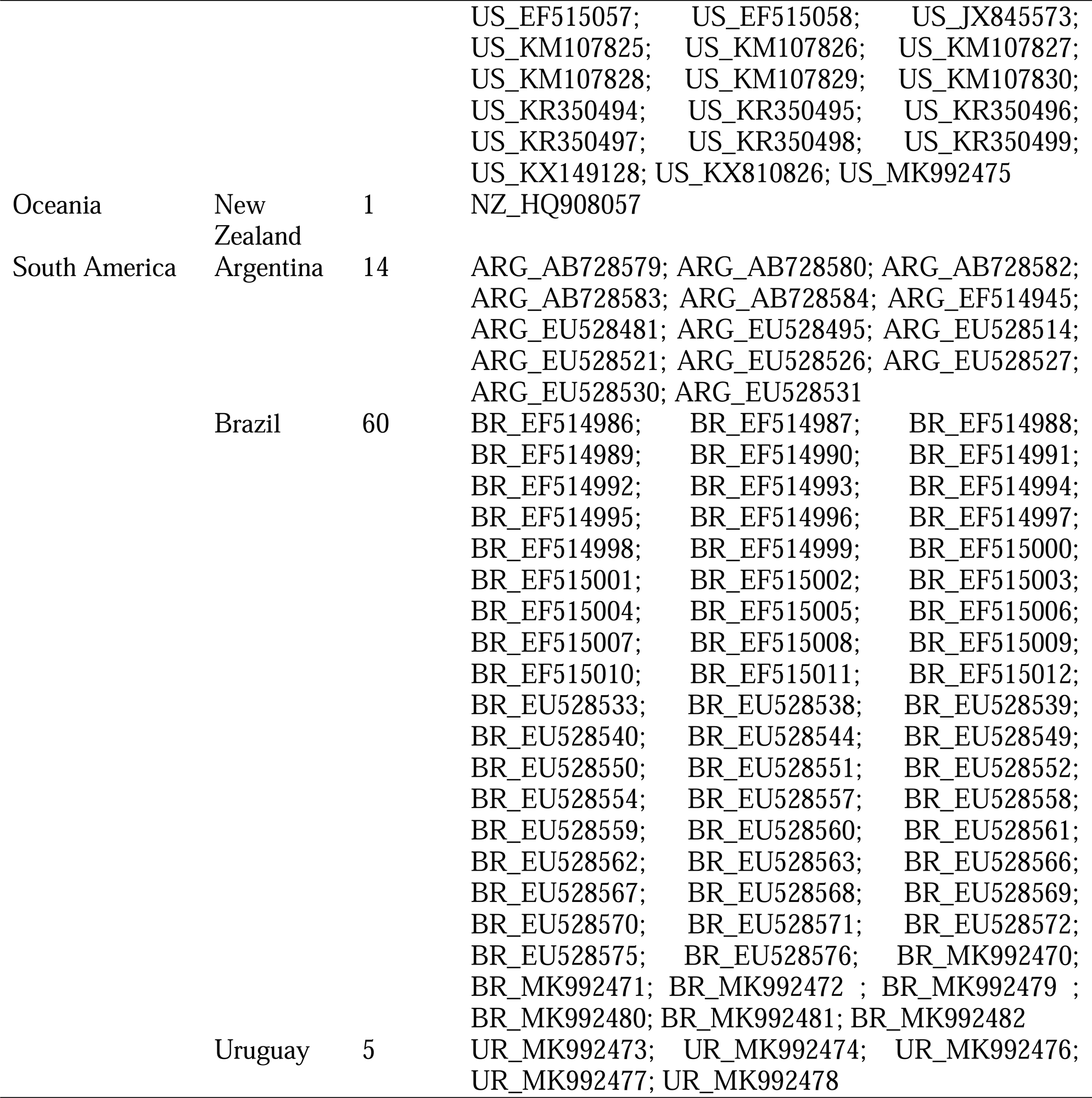
List of GenBank accession numbers used to prepare *Pomacea maculata* mitochondrial COI dataset (Dataset 2)

**APPENDIX 4.**
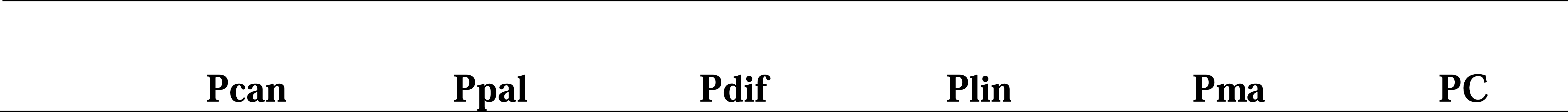
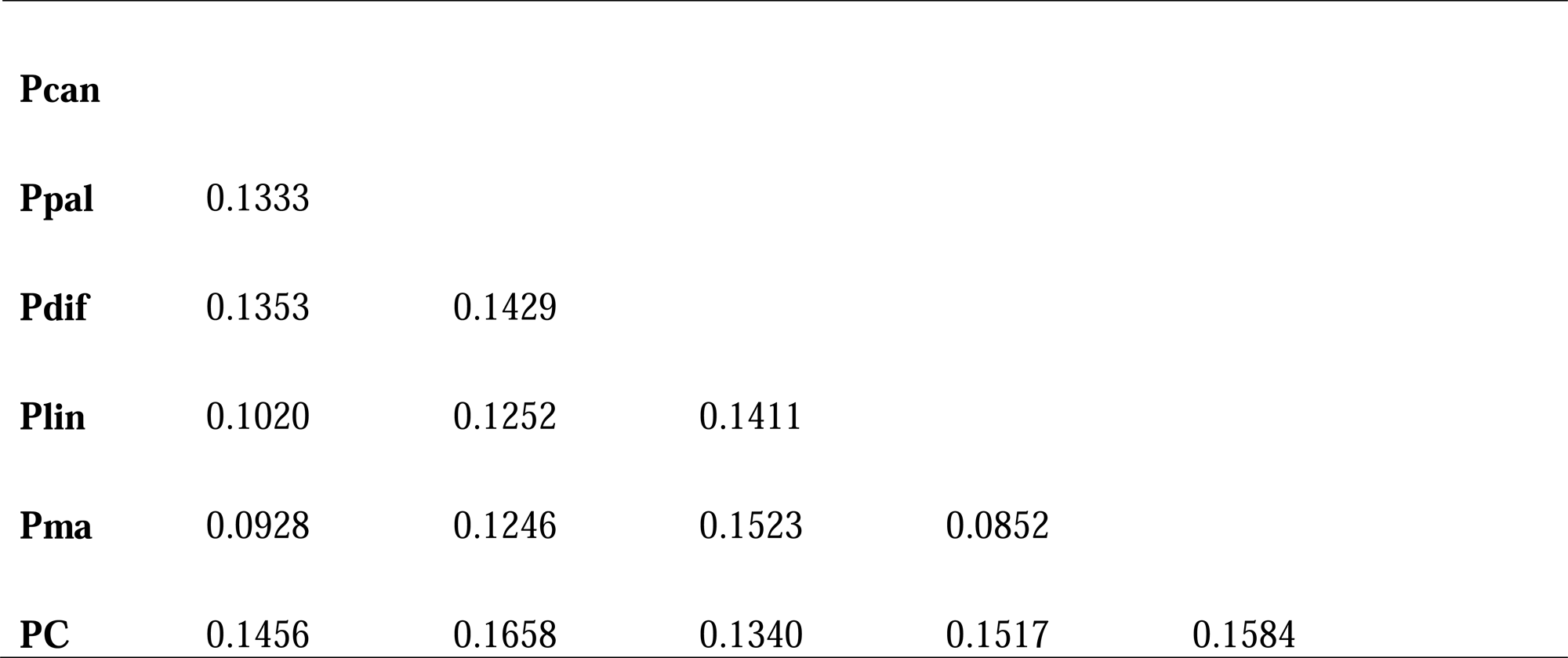
Genetic distance (p-distance) among the taxa used in the phylogenetic tree, Pcan:*P. canaliculate*; Pma: *P. maculata*; Plin: *P. lineata*, Ppal: *P. paludosa*, Pdif: *P. diffusa*, and PC: *P. scalaris*.

**APPENDIX 5.**
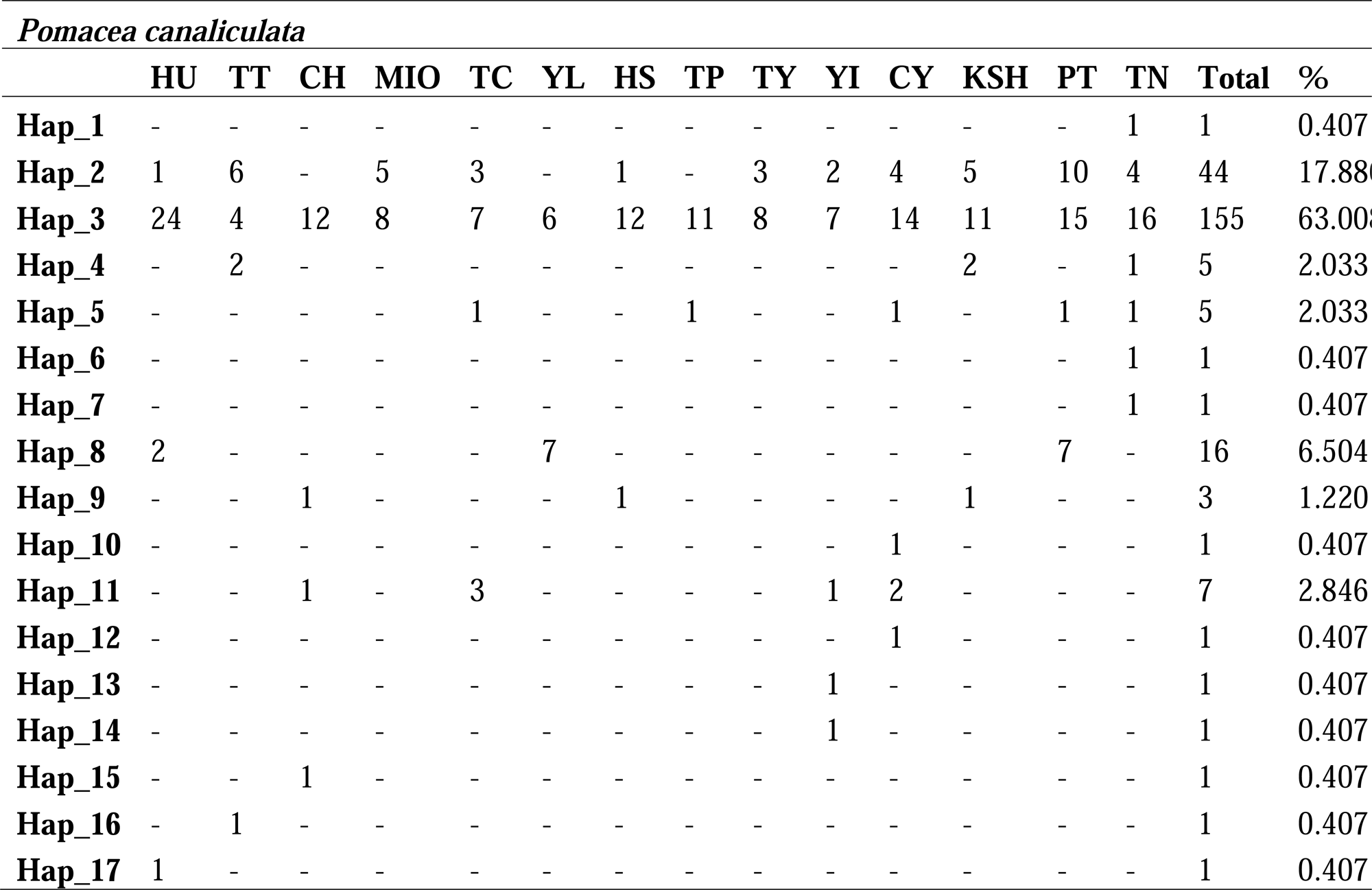
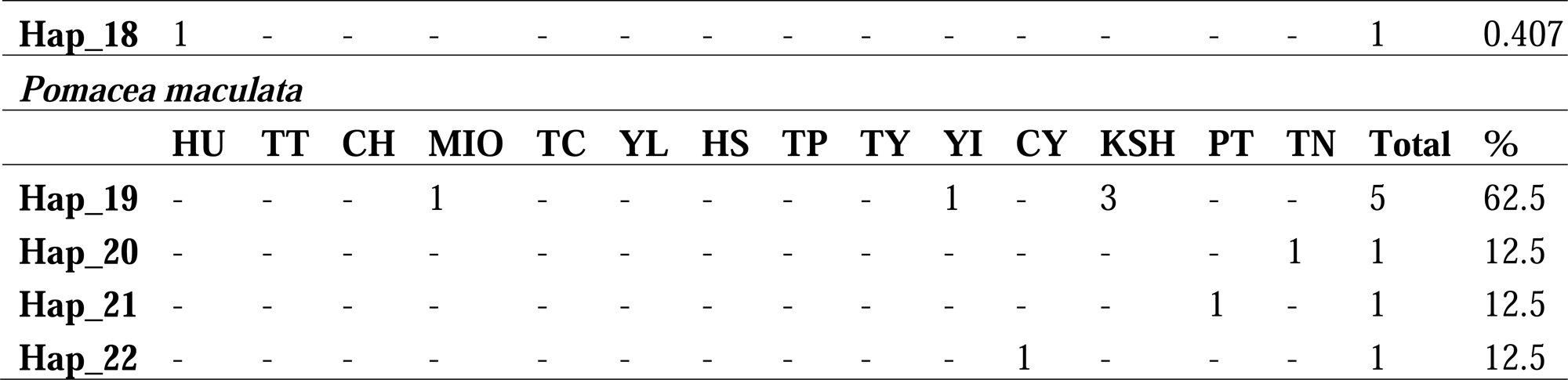
Haplotype distribution of *Pomacea* spp. in 14 counties/cities across Taiwan (full names of abbreviation can be found in APPENDIX 1.)

**APPENDIX 6.**
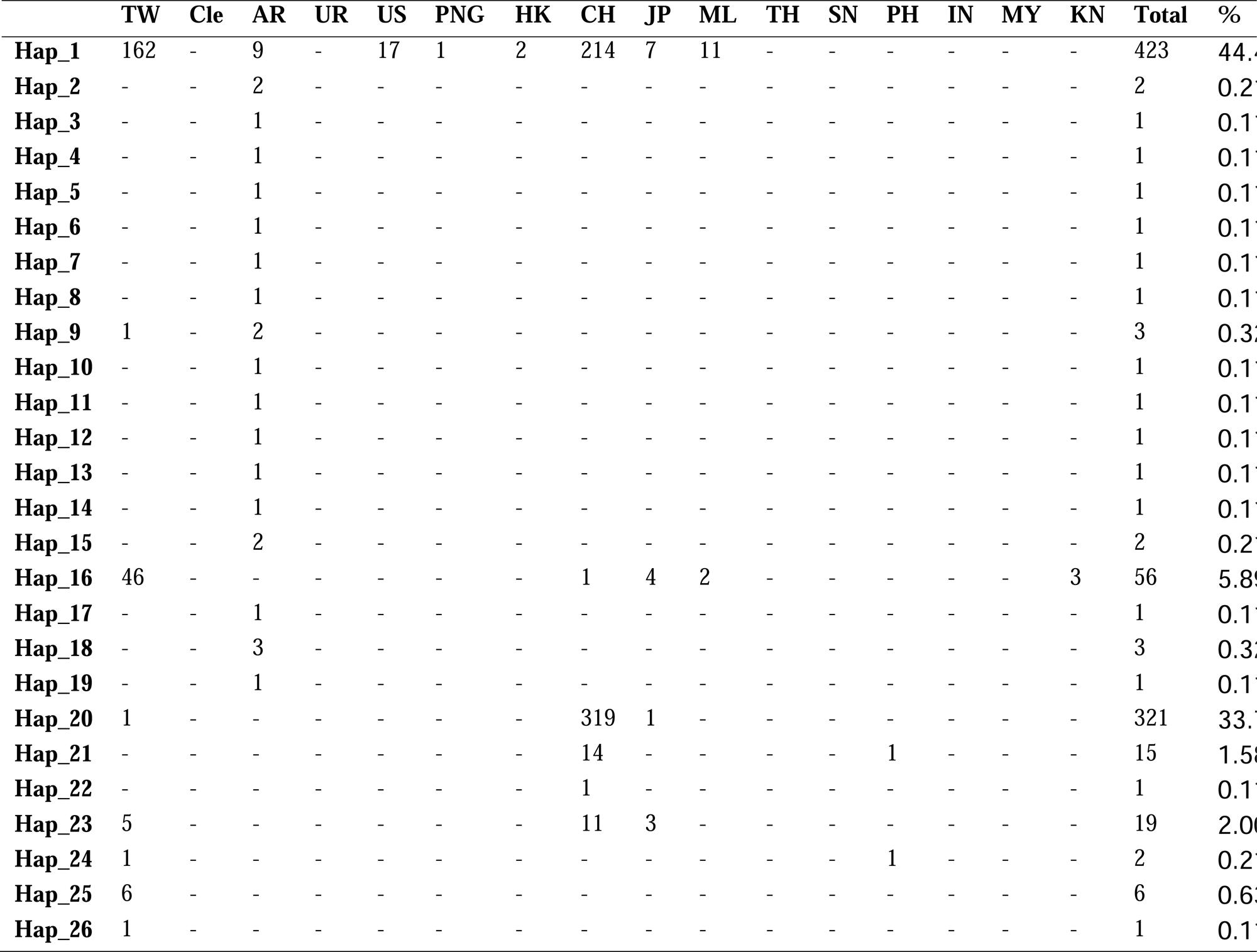
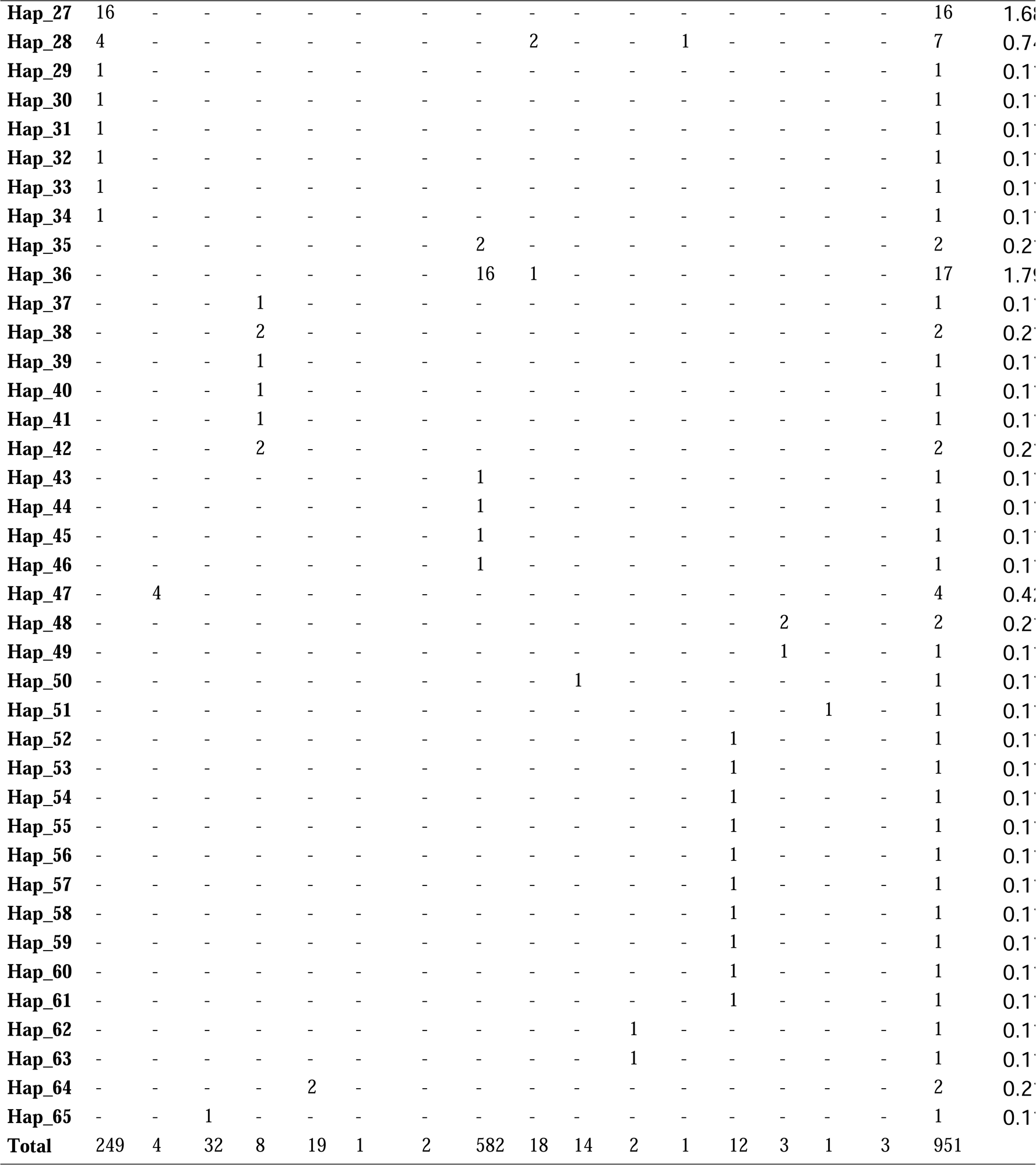
Haplotype distribution of *Pomacea canaliculata* from the global mitochondrial COI dataset (Dataset 3)

**APPENDIX 7.**
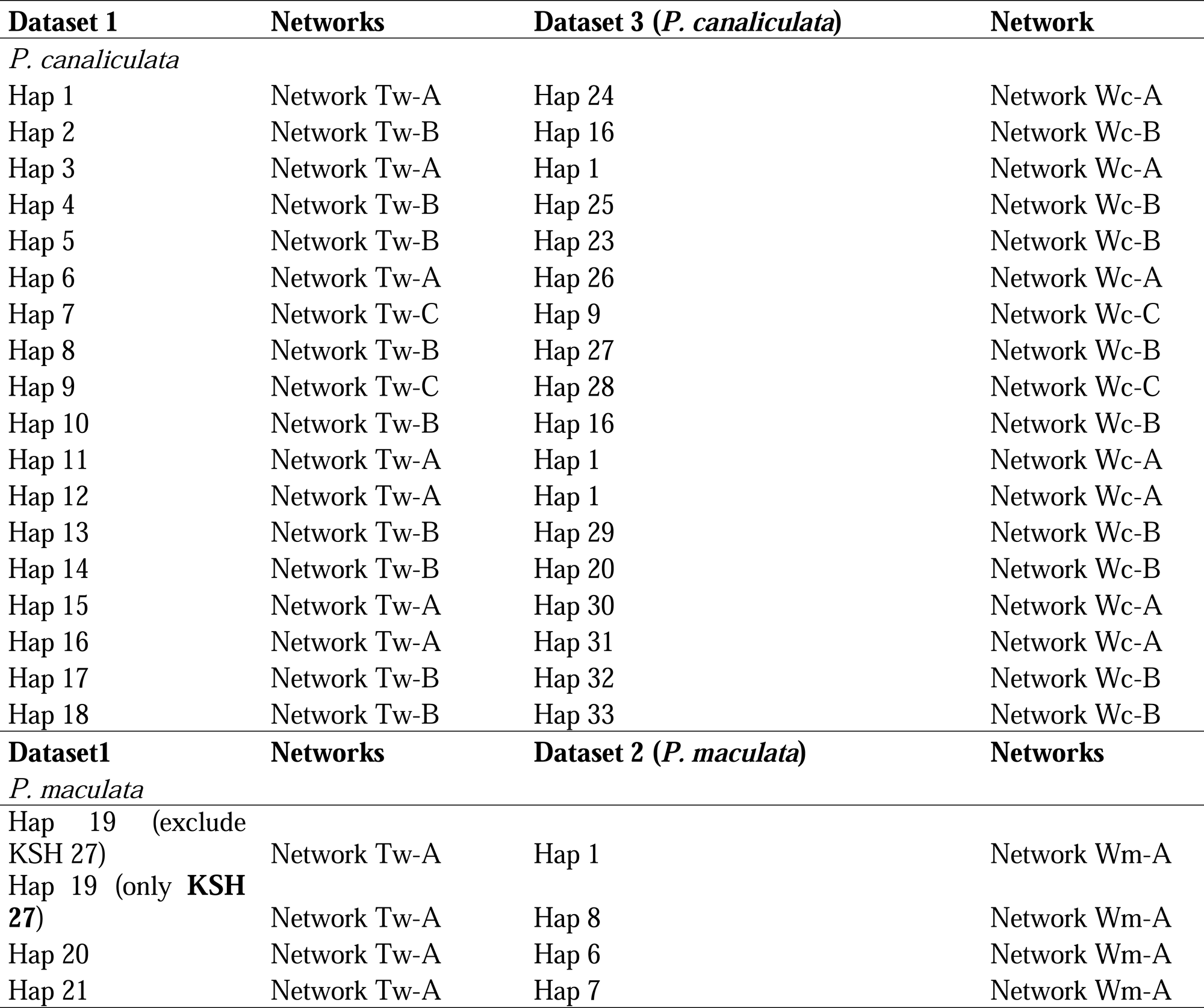
Comparison of Taiwan haplotype positions in different datasets, for *Pomacea canaliculata* Dataset 1 (Taiwan sequence) compared with Dataset 2 (global dataset of *Pomacea canaliculata*), and for *Pomacea maculata* Dataset 1 (Taiwan sequences) compared with Dataset 3 (global dataset of *Pomacea maculata*)

**APPENDIX 8.**
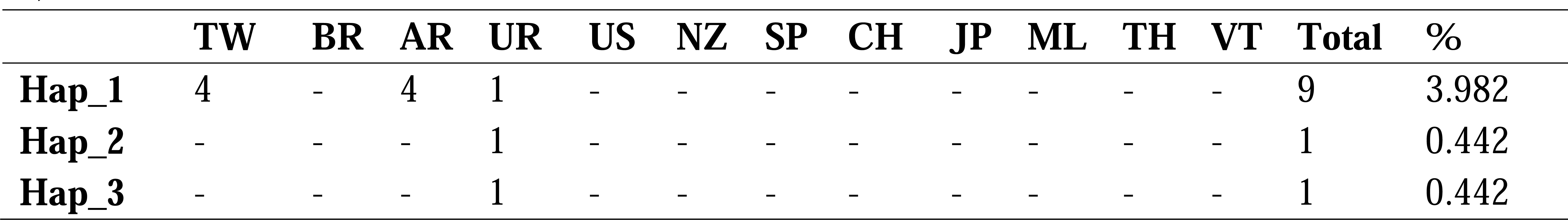
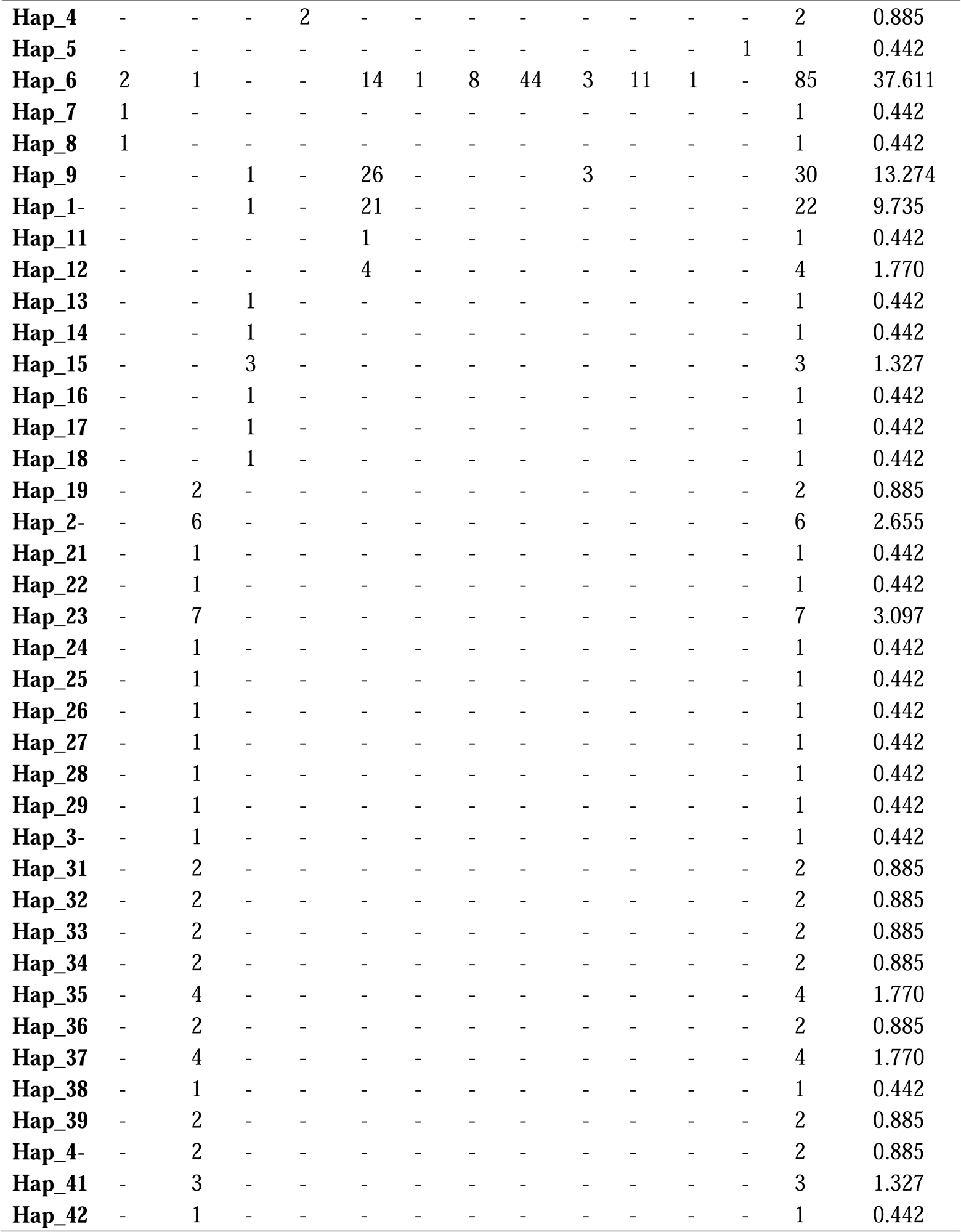
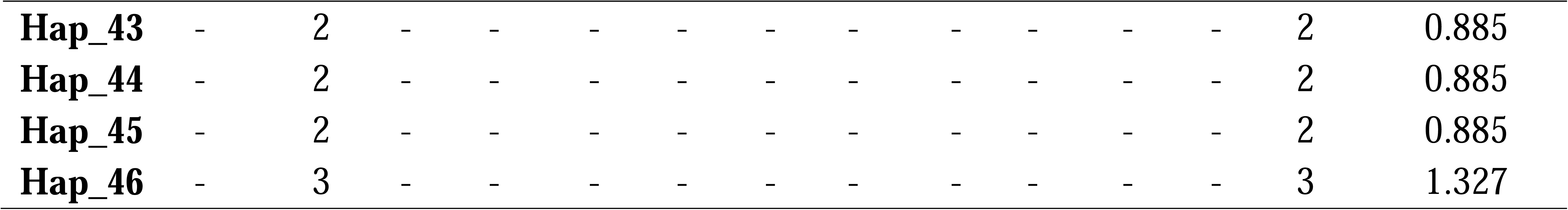
Haplotype distribution of *Pomacea maculata* from the global mitochondrial COI dataset (Dataset 2)

